# Hypersynchronous iPSC-derived *SHANK2* neuronal networks are rescued by mGluR5 agonism

**DOI:** 10.1101/2024.07.30.605451

**Authors:** Fraser P. McCready, Kartik S. Pradeepan, Milad Khaki, Wei Wei, Nicole Matusiak, Alina Piekna, Julio Martinez-Trujillo, James Ellis

## Abstract

Variants in the gene encoding the postsynaptic scaffolding protein SHANK2 are associated with several neurodevelopmental disorders, including autism spectrum disorder. Here, we used *in vitro* multielectrode arrays and pharmacological manipulations to characterize how functional connectivity and network-level firing properties were altered in cultures of human iPSC-derived *SHANK2* neurons. Using two isogenic pairs of *SHANK2* cell lines, we showed that the *SHANK2* hyperconnectivity phenotype was recapitulated at the network level. *SHANK2* networks displayed significantly increased frequency and reduced duration of network burst events relative to controls. *SHANK2* network activity was hypersynchronous, with increased functional correlation strength between recording channels. Analysis of intra-network burst firing dynamics revealed that spikes within *SHANK2* network bursts were organized into high-frequency trains, producing a distinctive network burst shape. Calcium-dependent events called reverberating super bursts (RSBs) were observed in control networks but rarely occurred in *SHANK2* networks. *SHANK2* network hypersynchrony and numbers of strong correlations were fully rescued by the group 1 mGluR agonist DHPG, that also restored detection of RSBs and significantly improved network burst frequency and duration metrics. Our results demonstrate that *SHANK2* variants produce a functional hyperconnectivity phenotype that deviates from the developmental trajectory of isogenic control networks. Furthermore, the hypersynchronous phenotype was rescued by pharmacologically regulating glutamatergic neurotransmission.

## INTRODUCTION

Autism spectrum disorder (ASD) is associated with protein coding and regulatory variants of genes that are important for proper synaptic function (Trost et al., 2022, Satterstrom et al., 2020, Wamsley et al., 2024). *SHANK2* is a high-confidence ASD risk gene with heterozygous loss-of-function variants being associated with ASD in humans (Berkel et al., 2010, Berkel et al., 2012, Leblond et al., 2012). The SH3 and multiple ankyrin repeat domains (*SHANK*) gene family (Monteiro and Feng, 2017, Leblond et al., 2014) encodes multidomain scaffold proteins that primarily function at the postsynaptic density (PSD) of excitatory synapses and are regulated by zinc (Daini et al., 2021, Vyas et al., 2021, Ha et al., 2018, Eltokhi et al., 2021). SHANK scaffolds anchor neurotransmitter receptors including NMDARs and AMPARs into complexes together with PSD95 and POSH (Yao et al., 2022, Shi et al., 2017). SHANKs also interact with HOMER proteins to couple group 1 metabotropic glutamate receptors (mGluR) in a signalling complex (Tu et al., 1999, Hayashi et al., 2009, Scheefhals et al., 2019). mGluR signalling leads to increased translation of synaptic proteins (Santini and Klann, 2014) that modulates synaptic plasticity (Waung and Huber, 2009) influencing the development of neural circuitry. Dysregulation of this process can lead to the formation of hyperconnected or hypoconnected neurons in neuropsychiatric disorders (Citri and Malenka, 2008). Altered synaptic plasticity is observed in *Shank2* mice (Schmeisser et al., 2012, Won et al., 2012, Vyas et al., 2021) although distinct phenotypes observed in homozygous mice are dependent on the *SHANK2* variant tested (Eltokhi et al., 2018, Lee et al., 2024).

One way to model heterozygous *SHANK2* variants is to isolate human induced pluripotent stem cells (iPSC) from ASD subjects and differentiate them into cortical neurons. We previously showed that iPSC-derived neurons harboring ASD subject-specific variants in *SHANK2* are hyperconnected, having increased dendrite length and complexity and making more synaptic connections with other cells in culture (Zaslavsky et al., 2019). In addition, *SHANK2* neurons had increased excitatory synaptic function including increases in spontaneous excitatory postsynaptic potentials (sEPSC) frequency and amplitude in intracellular recordings. Chronic treatment of *SHANK2* neurons with the mGluR agonist (S)-3,5-Dihydroxyphenylglycine (DHPG) prevented the increased dendrite length phenotype in neurons carrying a heterozygous nonsense variant (*SHANK2* R841X). DHPG was also used to demonstrate that Shank proteins are required for mGluR5 internalization and signalling in mouse neurons (Scheefhals et al., 2019, Verpelli et al., 2011). Further evidence supporting a role for dysregulated mGluR signaling in *SHANK2* ASD is that mGluR5 protein was reduced in cultured iPSC-derived *SHANK2* subject neurons and in the striatum of P7 *Shank2*(+/-) and *Shank2* (-/-) mice (Lutz et al., 2021). In addition, this reduced abundance of mGluR5 was accompanied by dysregulation of the extracellular signal-regulated kinase 1/2 (ERK1/2) signaling pathway, which is known to function downstream of mGluR5 activation.

*In vitro* multielectrode array (MEA) circuitry studies on iPSC-derived neurons often complement single neuron patch-clamp results (Deneault et al., 2019, Mok et al., 2022, Frega et al., 2019, Pradeepan et al., 2024). They allow longitudinal recordings of connectivity, and track the establishment of functional network burst events or emergence of more complex patterns of network burst activity that reverberate (Lau and Bi, 2005, Volman et al., 2007). Such reverberating super bursts (RSBs) have been found in typical and neurodevelopmental disorder networks *in vitro* (Pradeepan et al., 2024, Doorn et al., 2024). While our previous findings show that *SHANK2* neurons are hyperconnected at the single neuron level, it is unknown how this increased synaptic connectivity and glutamatergic signaling impact network activity dynamics. DHPG treatment for 24 hours was shown to enhance spike frequency of mouse neuronal networks in MEAs (Liu et al., 2020). However, the impact of DHPG treatment on the functional activity of *SHANK2* neurons remains entirely unexplored.

To address these questions, we utilized *in vitro* MEAs to extend the functional characterization of iPSC-derived *SHANK2* neurons to their network-level firing activity and collective firing dynamics. MEA recordings on isogenic pairs of iPSC-derived homozygous (-/-) *SHANK2* knockout (KO) and heterozygous (+/-) *SHANK2* R841X networks *in vitro* were analyzed for their patterns of activity from week 4 to week 8 of development. These *SHANK2* networks were hyperactive and hypersynchronous compared to controls, displaying network bursts that were significantly more frequent and shorter in duration. Calcium-dependent RSBs were detected in the isogenic control networks but were almost undetectable in *SHANK2* networks. Finally, chronic treatment of *SHANK2* networks with DHPG rescued network synchrony, and significantly improved the network burst frequency and duration, as well as partially restoring the proportion of RSBs. Overall, the hypersynchronous *SHANK2* network phenotype deviated from the developmental trajectory of isogenic controls, and was rescued by pharmacologically regulating glutamatergic neurotransmission.

## MATERIALS AND METHODS

### iPSC culturing and maintenance

IPS cells were cultured on Matrigel (Corning) or Geltrex (Gibco) coated plates with 1.5-2 ml of mTeSR1 media (STEMCELL Technologies), media were changed daily except the day after passing. IPS cells were passed once a week with ReLeSR (STEMCELL technologies) into 1:3-1:10 dilution depending on the density of the culture. Accutase (InnovativeCellTechnologies) was used for single-cell dissociation while cells cultured in MTeSR1 media supplemented with 10 μM Rho-associated Kinase(ROCK) inhibitor. Mycoplasma test was performed routinely.

### Ngn2 neuronal differentiation

Ngn2 induced cortical neurons were differentiated as previously described (Zhang et al., 2013, Mok et al., 2022). Briefly, On day 0 IPS cells were dissociated into single cells with accutase, 300,000 - 750,000 cells in 2 ml of mTeSR1 Supplemented with 10 μM ROCK inhibitor were seed into each well of Matrigel-coated 6 well plates. On Day 1 and day 2 cells were cultured with CM1 media [DMEM-F12(Giboco), 1 x N2 (Gibco), 1 x pen/strep(Gibco), 1 x NEAA(Gibco), 1 μg/ml laminin (sigma)] supplemented with BDNF (Peprotech) and 10 ng/mL GDNF (Peprotech) and 2 μg/mL doxycycline hyclate (Sigma) with the exception that day 1 cells were still cultured in 10 μM Rock inhibitor and on day 2, Rock inhibitor was withdrawn and cell selection was started with (2–5 μg/mL) puromycin(Sigma). From day 3 to day 8, cells were cultured in CM2 media [Neurobasal media (Gibco), 1 x B27(Gibco), 1 x Glutamax (Gibco), 1 x pen/strep, 1 μg/ml laminin 10 ng/ml BDNF, 10 ng/ml GDNF] supplemented with 2 μg/mL doxycycline hyclate, where on day 3 the cells were still under puromycin selection and from day 4 to day 8, the selection was withdrawn. From day 6-8, 10 μM Ara-C (Sigma) was added to CM2 media. On day 8, post-Ngn2 induced neurons were dissociated with accutase and filtered through a 70m filter reseeded on plates for future assays. *FUW-TetO-Ngn2-P2A-EGFP-T2A-puromycin* and *FUW-rtTA* plasmids for excitatory cortical neuron differentiations were kindly gifted by T. Sudhof (Zhang et al., 2013).

### MEA plating and recording

Day 8 post-*Ngn2* induced neurons were plated on MEA plate as previously described (Mok et al., 2022). Briefly, 12 well MEA plates with 64 electrodes (Axion Biosystems) were coated with filter sterilized 0.1% PEI solution in borate buffer pH 8.4 at room temperature for 1 hour, followed by 4 times wash with water and dried overnight. 100,000 of Day 8 *Ngn2* neurons in 100μl of droplets were seeded on each well in CM2 Brainphys media [Brainphys (STEMCELL Technologies), 1 x Glutamax, 1 x pen/strep 10 ng/ml BDNF, 10 ng/ml GDNF] supplemented with 400 μg/ml laminin and 10 μM ROCK inhibitor. Cells adhered to the well with hydration for 2 hours and each well was topped up with 1 ml of CM2 brainphys media supplemented with 40μg/ml laminin. Next day 20,000 P1 mouse astrocytes were added to each well. Media was changed twice a week exactly 24 hours before recording.

For MEA recordings, each plate was allowed to incubate for 5 minutes on a Axion Maestro device heated to 37°C under 5% CO_2_. Spontaneous activity was then recorded at a sampling frequency of 12.5 kHz for 5 minutes using AxIS v2.0 software (Axion Biosystems), with the analog settings set to “Field Potentials” recording mode (1200X gain, 1 – 2000 Hz bandwidth, median referencing). The analog settings were then changed to “Neural: Spikes” mode (1200X gain, 200 – 5000 Hz bandwidth, median referencing) and an additional 5 minutes of spontaneous activity was recorded. Field potential recordings were used exclusively for correlated spectral entropy synchrony analysis (see associated methods section). Neural spikes recordings were further bandpass filtered at 0.2 – 3 kHz and spikes were detected using a threshold crossing method with the threshold set at 6x the standard deviation of the noise of recording electrodes. These data were used as the starting point for all other network analyses performed.

### Network burst detection and offline analysis

Network burst detection and offline analysis of MEA recordings was performed using Neural Metric Tool v2.5.7 software (Axion Biosystems). Electrodes were considered active if spikes were detected at a rate of at least 5 spikes per minute. Single-channel bursts were detected using the poisson surprise algorithm with the minimum surprise parameter set to 3. Network burst detection settings were tailored for each individual well to avoid spurious network burst calls and can be found listed in Supplemental Table 1. Supplemental neural metrics and network burst lists were exported for each recording and further analyses were completed in RStudio, Python, and MATLAB.

### Correlated spectral entropy synchrony analysis

Correlated spectral entropy based network synchrony analysis was performed as described (Kapucu et al., 2016). Field potential recordings were first filtered with a 60 Hz notch to remove powerline noise, then a 7 HZ high pass filter to remove low-frequency fluctuations. The filtered voltage signal for each electrode was then split into 0.5 second windows, with 50% overlap between consecutive windows. The frequency power spectrum of each segment, *P*(*f*), was calculated using the *pwelch* function in MATLAB R2022a and was normalized by the total power of each segment:

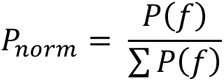

The spectral entropy of each signal segment (S_i_) was then calculated as

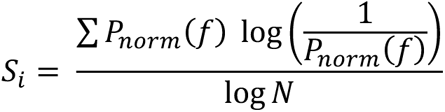

Where ∑ *P_norm_*(*f*) is the sum of the normalized power spectrum for a signal segment containing *N* samples (e.g. for our recordings, *N* = 12500 for a 1 second segment of an extracellular voltage signal recorded at a sampling frequency of 12.5 kHz).

The cross-correlation of the spectral entropy of two signals, *S_x_* and *S_x_*, at lag *l* = 0 was then estimated using the *crosscor* function in MATLAB R2022a. This cross-correlation values is reported as the CorSE synchrony score for each electrode pair.

### Average network burst alignment

For each recording, intra-network burst spike times were binned in 10 ms time intervals. Binned network bursts were then zero-padded and aligned in a window ranging from t = -3 seconds to t = 3 seconds, with the bin containing the greatest number of spikes centered at t = 0 (representing the “burst peak”). The average network burst for each recording was then estimated by calculating the average number of spikes per bin across all aligned network bursts from that recording. A final “average network burst heatmap” for each cell line were then generated by plotting the number of spikes per bin in the averaged network burst from each recording.

### Formulation of the AceDimer Machine Learning Algorithm

The AceDimer algorithm core function is to evaluate feature contributions in neuroinformatic datasets. This involves generating feature subsets and assessing their impact on classification accuracy. The process can be summarized as follows:

1. Feature Subset Generation: AceDimer generates all possible subsets of features for a dataset with N features. Each subset *S_i_* is a combination of features where *i* is a subset of (1, 2, …, N).

a. S_i_ = (f1, f2, …, fm), where m ≤ N
2. Classification Accuracy Calculation: The classification accuracy A(Si) is calculated using the chosen classifier for each feature subset Si.

_b._ A(S_i_) = Accuracy of Classifier with Features S_i_
3. Feature Contribution Analysis: The contribution of each feature is assessed by comparing the accuracy of models with and without the feature. For a given feature f_j_, its contribution C(f_j_) is determined by the change in accuracy when fj is included in the subset.

C(f_j_) = A(S_i_ ∪ (fj)) - A(S_i_ \ (fj))
4. Overall Feature Importance: The overall importance of each feature is calculated by taking the sum of its contributions across all subsets where it appears and then dividing this sum by the standard deviation of the feature’s contributions across these subsets. This method offers a normalized measure of a feature’s importance, accounting for the variability of its contributions.

The formula for the overall importance of a feature f_j_ is as follows:

OI(f_j_) = (Sum of C(f_j_) for all *i* where f_j_ is in S_i_) / Standard Deviation of C(f_j_) Where:

- “Sum of C(f_j_) for all *i* where f_j_ is in S_i_” represents the total contribution of the feature f_j_ across all subsets that include it.
- “Standard Deviation of C(f_j_)” refers to the standard deviation of the contributions of the feature f_j_ across these subsets.

This updated method in AceDimer offers a more detailed and balanced analysis of each feature’s significance, considering its contribution and its consistency across different combinations.

These steps and calculations form the basis of the AceDimer algorithm, allowing for a comprehensive analysis of feature contributions and their interdependencies in neuroinformatic data.

### Pharmacological treatments

All MEA plates were recorded twice per week, using the same recording practices and filter settings described in the MEA Plating and Recording section. During each recording, all wells were monitored for the appearance of RSBs through simple visual inspection of live raster plots generated by on-line spike and network burst detection in AxIS v2.0 software. Pharmacological treatments began after RSBs were detected in cultures. Before testing of pharmacological compounds, cell culture media was changed with fresh CM2 media and plates were allowed to incubate for 1 hour to allow neuronal activity to stabilize. A baseline recording of spontaneous neural activity was then taken for 10 minutes at 37 °C. Immediately following baseline recording, one pharmacological agent (0.1% DMSO, or 10 µM bicuculline) was added to each well, and an additional 10 minutes of spontaneous activity was recorded in the presence of drug compound. Washout of pharmacological agents was then achieved by changing culture media with fresh CM2 a total of 3 times, and plates were then allowed to incubate for another hour to allow activity to stabilize. A new 10-minute baseline recording was then taken, and the process was repeated until all pharmacological agents had been tested. 25 µM EGTA-AM was always added as the final drug compound in the rounds of testing as it was observed that normal baseline spontaneous activity did not return after drug washout.

### Statistical analysis

Statistical tests were performed in RStudio v2023.06.0 running R v4.3.1. Normality assumptions were formally tested using a Kolmogorov-Smirnov normality test. Comparisons between isogenic pairs were conducted by Mann-Whitney test with Benjamini-Hochberg correction for multiple testing (*p < 0.05, **p < 0.01, ***p < 0.001).

### Data visualization

Representative raster plots were generated in Python using the eventplot function from the Matplotlib library (v3.7.1). Representative voltage races were plotted in MATLAB R2022A using functions provided by Axion Biosystems. Circular connectivity plots were generated in Python using the circular_layout and plot_connectivity_circle functions from the MNE tools library (v1.4.2). CorSE heatmaps were generated in Python using the heatmap function from the seaborn library (v0.12.2). CorSE network maps were generated in RStudio using the graph_from_data_frame function from the igraph package (v1.4.3). All other plots were generated in Rstudio using the ggplot function from the ggplot2 package (v3.4.2). Margin distributions were added to 2D scatterplots using the ggMarginal function from the ggExtra package (v.0.10.0).

## RESULTS

### iPSC-derived SHANK2 neurons assemble into functional networks in vitro

We previously reported that ASD-associated variants in *SHANK2* result in hyperconnectivity of single neurons. To examine how population-level network activity is impacted by *SHANK2* variants, we used *in vitro* MEAs to record spontaneous firing activity from *SHANK2* neurons. We used two pairs of isogenic *SHANK2* and control lines that were previously described (Zaslavsky et al., 2019) (Figure 1A). The first isogenic pair consists of an ASD subject-derived line which harbours a single base pair non-sense variant (*SHANK2* R841X) and an isogenic correction line (R841X-C), which was generated by using CRISPR-Cas9n gene editing to correct the point mutation back to the wild-type allele. The second pair consists of a control line reprogrammed from an unaffected control individual (CTRL) and an isogenic, homozygous knockout line (*SHANK2* KO).

**Figure 1.**
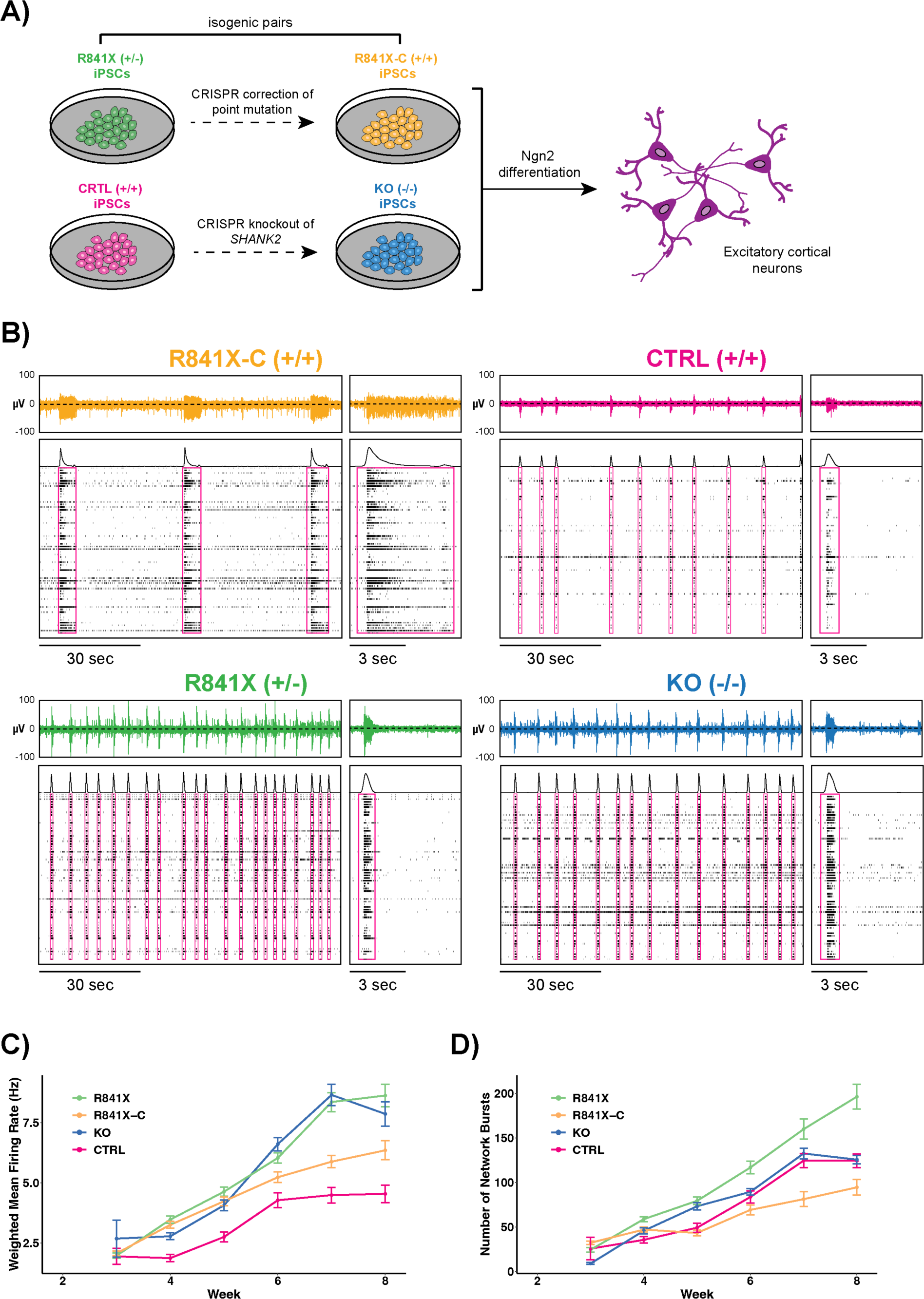
iPSC-derived *SHANK2* neurons assemble into functional networks *in vitro.* (**A**) Overview of isogenic iPSC lines and workflow to generate excitatory cortical neurons used in this study. (**B**) Representative extracellular voltage signals and raster plots of isogenic *SHANK2* and control neuron networks taken from recordings at week 7 of development. (**C**) Weighted mean firing rate and (**D**) the number of detected network bursts were observed to increase over time for all lines.

Excitatory neurons were generated from each line by forced overexpression of the pro-neurogenic transcription factor *Neurogenin2* (*Ngn2*) and were subsequently co-cultured with primary mouse astrocytes on 12-well multielectrode array plates (Axion Biosystems) containing 64 extracellular electrodes per well. Five minutes of spontaneous network activity was recorded twice a week, starting from 16 days after *Ngn2* induction (DIV16) until DIV60. All four lines developed robust spontaneous firing activity, with the weighted mean firing rate (wMFR) of all lines increasing from week 2 – week 8 of recording (Figure 1B-C). Additionally, we observed network bursts by week 3 in all lines, with the average number of detected network bursts increasing from week 3 – week 8 (Figure 1D). Network bursts are a stereotyped pattern of spontaneous activity where bursts of high-frequency spiking activity are detected across multiple spatial locations in the network simultaneously, and their appearance is indicative of the establishment of network circuitry and functional network activity. These results indicate that neurons derived from all four iPSC lines produce robust spontaneous spiking activity and self-organize into functional networks.

### Optimization of network burst detection

Our detection of network burst events in the MEA recordings was initially handled by automated network burst detection algorithms. Importantly, the choice of detection algorithm and its associated parameters must be optimized to ensure that network burst identification is accurate. Indeed, suboptimal detection has been shown to significantly influence the results of downstream network activity analyses and phenotyping metrics (Mossink et al., 2022). Thus, to ensure the reliability of our network metrics, we ran automated network burst detection using *fixed* interspike interval (ISI) threshold, *adaptive* ISI threshold, and *envelope detection* algorithms (see methods for a detailed description of each detection algorithm). We then manually inspected the resultant network burst calls for every recording to evaluate the accuracy of each algorithm. We found that no single set of algorithm and parameter settings was appropriate for accurate network burst detection in all cultures across all time points, and this bulk approach to detection led to erroneous network burst calls in a significant number of recordings (Supplemental Figure 1). We found that ISI threshold algorithms tended to underestimate network burst frequency and overestimate network burst duration in many of our recordings grouping many distinct network bursts together as a single event. Accordingly, we instead chose to tailor network burst detection parameters for each culture individually. The detection algorithm and parameter settings used for each recording are listed in Supplemental Table 1.

### SHANK2 neurons collectively fire frequent network bursts with short durations

To begin characterizing population-level differences in network firing dynamics, we first requantified the mean firing rate (MFR) in recordings of *SHANK2* and isogenic control networks from week 4 (DIV29 – DIV35) to week 8 (DIV57 – DIV63) of development. We found that *SHANK2* networks were significantly more active than their respective isogenic controls, with the *SHANK2* R841X and *SHANK2* KO lines both exhibiting a 1.8-fold and 2-fold increase in MFR at week 7, respectively (Figure 2A). The wMFR was also increased in *SHANK2* networks, indicating that the increased firing rate was not caused by an increase in the number of active recording sites in *SHANK2* cultures (Figure 2B). In addition to an increase in baseline firing rate, *SHANK2* networks displayed a significant increase in the frequency of network burst events across all time points examined (Figure 2C). Average network burst frequency was increased 4.1-fold in *SHANK2* R841X cultures when compared to isogenic R841X-C networks at week 7 of recording. This was accompanied by a concurrent 2.6-fold increase in the frequency of single-channel bursts (Figure 2D). An increased frequency of network bursting was also observed in *SHANK2* KO cultures (1.4-fold increase at week 7) (Figure 2C). A trend of increased single-channel burst frequency was also observed in *SHANK2* KO cultures, reaching statistical significance at week 4 and week 5 of development (Figure 2D).

**Figure 2.**
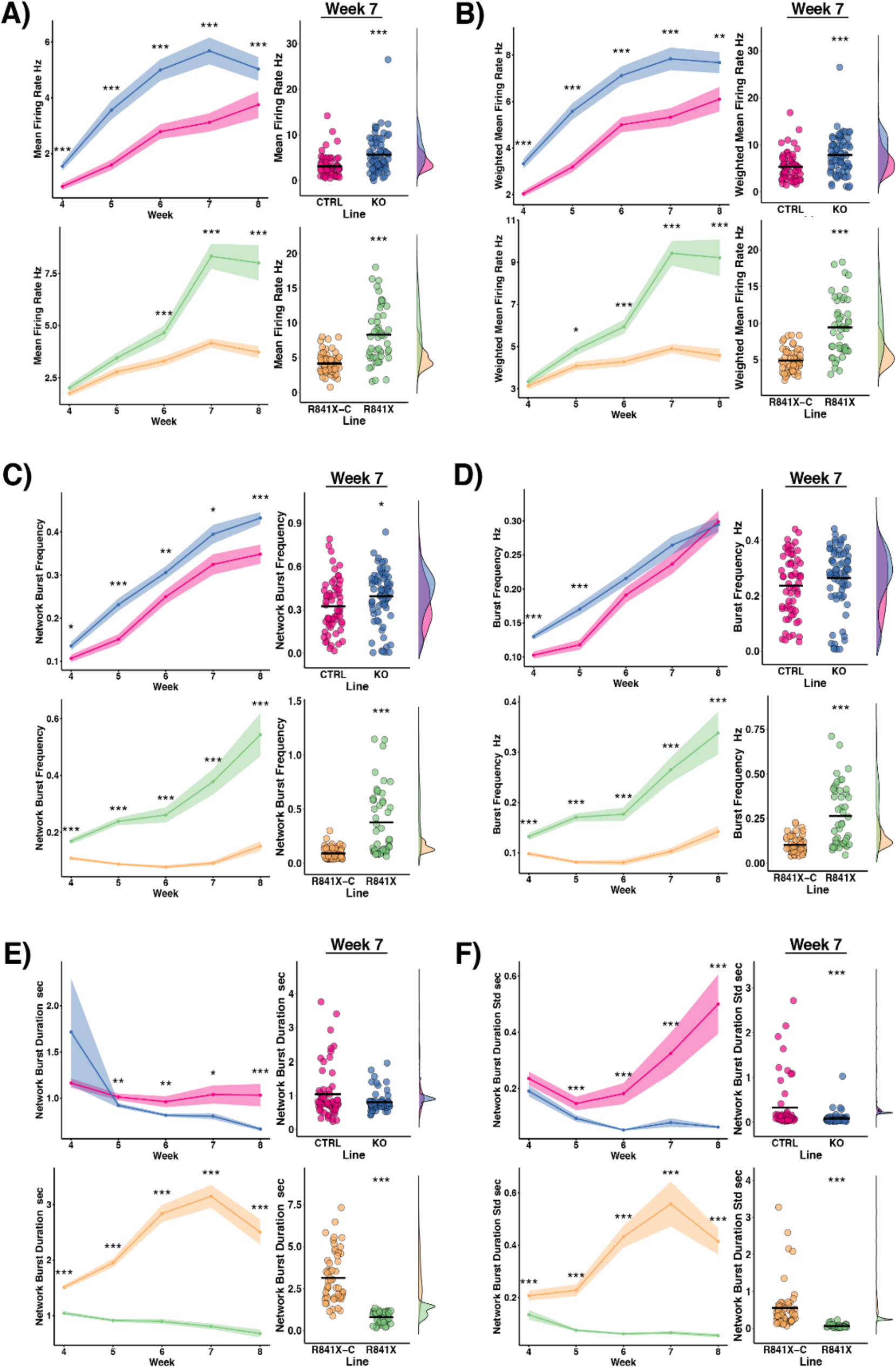
Increased network burst frequency and reduced network burst duration in *SHANK2* networks. (**A**) *SHANK2* KO (top) and *SHANK2* R841X (bottom) networks exhibit increased mean firing rate (MFR) when compared to their isogenic controls. (**B**) Normalizing MFR by the number of active recording channels does not change discrepancy in the wMFR observed in *SHANK2* networks. (**C**) Quantification of network burst frequency, (**D**) burst frequency, (**E**) network burst duration, and (**F**) the standard deviation of network burst durations in *SHANK2* and isogenic control networks. Shaded error bands on lineplots indicate mean ± SEM. Crossbars on dotplots indicate the mean, marginal density plots show distribution of datapoints (n = 54 for R841X-C, n = 47 for R841X, n = 60 for CTRL, n = 73 for KO. Network recordings were taken from 6 independent differentiations for each cell line). *P < 0.05, **P< 0.01, ***P < 0.005; Mann-Whitney U test with Benjamini-Hochberg correction for multiple testing.

Network bursts occurring in *SHANK2* cultures were considerably shorter in duration than those of control networks. Again, this difference was more pronounced in the ASD subject-derived *SHANK2* R841X networks, which showed a 3.9-fold reduction in average network burst duration compared to a 1.3-fold reduction in *SHANK2* KO networks (Figure 2E). In addition, the durations of network bursts within a given recording period were considerably less variable in *SHANK2* cultures than in controls. This resulted in a striking 9-fold reduction in the standard deviation of network burst durations in *SHANK2* R841X networks at week 7 of development, while a 4.3-fold reduction was observed for *SHANK2* KO cultures (Figure 2F). Together, these results indicate that *SHANK2* networks fire network bursts that are shorter, more regular in duration, and more frequent than controls.

### SHANK2 networks exhibit hypersynchronous firing activity

We next sought to characterize differences in network synchronization in *SHANK2* and control cultures by conducting a pairwise analysis of neural synchrony between all 64 recording channels in each network recording. To evaluate pairwise correlation strengths in our networks, we chose to employ the correlated spectral entropy (CorSE) method (Kapucu et al., 2016). In contrast to event-based synchrony measures which evaluate spike train synchrony by looking for correlations in the timing of binary spike events detected at different recording channels, the CorSE method takes the continuous voltage signal from each recording channel as inputs and evaluates synchrony by looking for correlated temporal changes in the contents of the signal’s frequency spectrum distribution (Figure 3A). A significant benefit to this approach is that it does not rely on spike detection prior to analysis and is thus not affected by sub-optimal spike detection, which could arise from high signal noise, low-amplitude spikes, or experimenter choice of detection algorithm. In addition, since the CorSE method is performed on continuous filtered voltage signals, low-frequency information contained in local field potentials (LFPs) generated by subthreshold and population-level activity is retained in the analysis and contributes to synchrony calculations (Kapucu et al., 2016).

**Figure 3.**
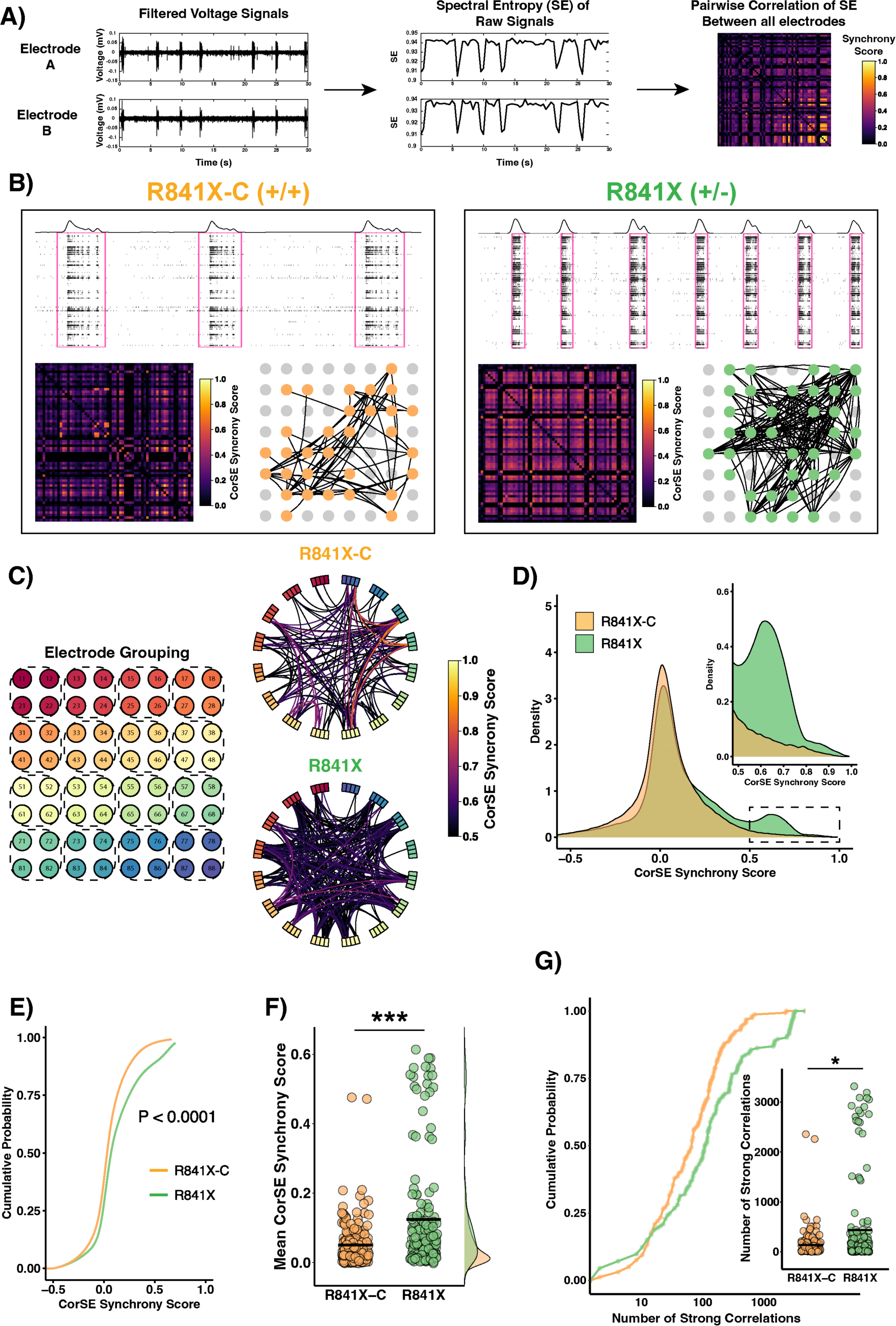
Increased synchrony of *SHANK2* R841X networks as measured by correlated spectral entropy. (**A**) Approach for calculation of synchrony pairwise correlations based on correlated changes in the spectral entropy of raw voltage signals. This method does not require spike detection. (**B**) Representative CorSE output for *SHANK2* R841X and R841X-C networks. Each sub-panel shows a raster plot of 15 seconds activity from the 5-minute recordings and corresponding correlation matrix showing the synchrony score for each electrode pair (bottom left). The bottom right panel shows the 8 x8 electrode grid with an overlaid a connectivity map for the recording. Each edge represents a connection between two electrodes with a synchrony score > 0.5. Coloured nodes indicate electrodes involved in these connections and are shown at their appropriate spatial location on the electrode grid. (**C**) Circular connectivity plots from representative recordings showing all pairwise correlations with a strength > 0.5. Boxes around the circumference of the circle represent electrodes and edges represent connections. Edge colour indicates the strength of connection. Electrode grouping schematic shows how electrodes in the 8 x 8 electrode grid are grouped in circular connectivity plots. (**D**) Smoothed gaussian kernel density estimates comparing the distribution of connection strengths across all recordings at week 7 and 8 in *SHANK2* R841X and R841X-C networks. (**E**) Empirical cumulative distribution functions comparing CorSE synchrony scores in R841X-C and *SHANK2* R841X networks. (**F**) Quantification of mean synchrony scores in all recorded networks. (**G**) *SHANK2* networks show increased number of strong correlations than controls (n = 100 for R841X-C, n = 80 for R841X. Network recordings were taken from 6 independent differentiations for each cell line). *P < 0.05, **P< 0.01, ***P < 0.005; single tailed two-sample Kolmogorov-Smirnov test for (E), Mann-Whitney U test for (F,G).

Accordingly, we also conducted network recordings with analog filter settings set to “field potential mode” (1 – 2000 Hz bandwidth, see Methods) to capture these low-frequency signal contributions and used these data for our functional connectivity analysis. Representative raster plots and corresponding adjacency matrices showing pairwise CorSE synchrony scores are shown in Figure 3B and Supplemental Figure 2A. Network graphs showing all pairwise connections with a synchrony score > 0.5 are also shown. These same data are also presented as circular connectivity graphs in Figure 3C to better visualize connections at the expense of retaining spatial information about the network structure.

To gain a broad understanding of how connectivity may be altered in established *SHANK2* networks, we pooled CorSE synchrony scores from mature *SHANK2* and control networks at weeks 7 and 8 of development and compared their distributions. We found that the distribution of correlation strengths from both *SHANK2* R841X and *SHANK2* KO networks showed a significant shift towards higher synchrony values in comparison to their isogenic controls (P < 0.0001, single tailed, two-sample Kolmogorov-Smirnov test, Figure 3D-E and Supplemental Figure 2B-C). Notably, while the distributions for all lines were approximately bell shaped with a primary peak centered around CorSE values close to zero, the *SHANK2* R841X distribution contained a prominent secondary peak centered around CorSE synchrony value of approximately 0.6, which was markedly absent from the other distributions (Figure 3D). While the *SHANK2* KO distribution also lacked this notable secondary peak at higher synchrony values, it contained greater density in its right tail than either control distribution (Supplemental Figure 2B).

To further quantify differences in pairwise correlation strengths between cell lines, we calculated the mean CorSE synchrony score for each network recording and compared the mean CorSE values from *SHANK2* and control cultures. As expected, mean synchrony scores were increased 3.1-fold in *SHANK2* R841X networks (Figure 3F) and 1.7-fold in *SHANK2* KO networks (Supplemental Figure 2D-E). Given the shift towards high CorSE synchrony values in *SHANK2* networks, we next asked how strong correlations develop in *SHANK2* cultures. As done previously (Kapucu et al., 2016), we defined strong correlations as any connection with a synchronization score greater than 0.5 and then quantified the number of strong correlations present in each network recording.

Concurrent with the increase in mean CorSE values, we found a significant 4.9-fold increase in the mean number of strong correlations in *SHANK2* R841X networks (Figure 3G) and a 2.2-fold increase in the number of strong correlations in *SHANK2* KO cultures (Supplemental Figure 2F). Taken together, these results show that *SHANK2* networks are hypersynchronous.

### Altered intra-network burst shapes and firing dynamics in SHANK2 networks

Given the shorter and more regular duration of network bursts in *SHANK2* cultures, we next asked how intra-network burst firing activity might be structured differently in *SHANK2* networks. A technique of network burst alignment and comparison of network burst shapes was used to characterize the effects of cadherin-13 knockdown on network activity in cultures of iPSC-derived neurons (Mossink et al., 2022). Moreover, such characteristic network burst shapes can be used to discriminate between healthy and disease state dopaminergic and motor neurons in an iPSC model of Parkinson’s disease (Ronchi et al., 2021). We noted that while our control network bursts generally appeared to consist of a mixture of rapid burst firing and tonic, low-frequency spiking, *SHANK2* network bursts appeared to consist primarily of high-frequency short burst firing (see Figure 1B). To better visualize these network burst shapes, we binned intra-network burst spikes from each recording into 50 ms time intervals and then calculated the average number of spikes per bin across all network bursts for each recording. Within each recording, detected network bursts were aligned so that the bin containing the greatest number of spikes (i.e. the network burst “peak”) was centered at time t = 0 before averaging. Intra-network burst spiking intensity was then visualized by plotting this aligned, average network burst from each recording together to produce an average network burst heatmap for each line (Figure 4A). Moreover, by plotting the average number of spikes per bin across each heatmap, we were able to obtain a characteristic network burst shape for each cell line, which provides insight into how intra-network burst firing rates change as a function of time (Figure 4B).

**Figure 4.**
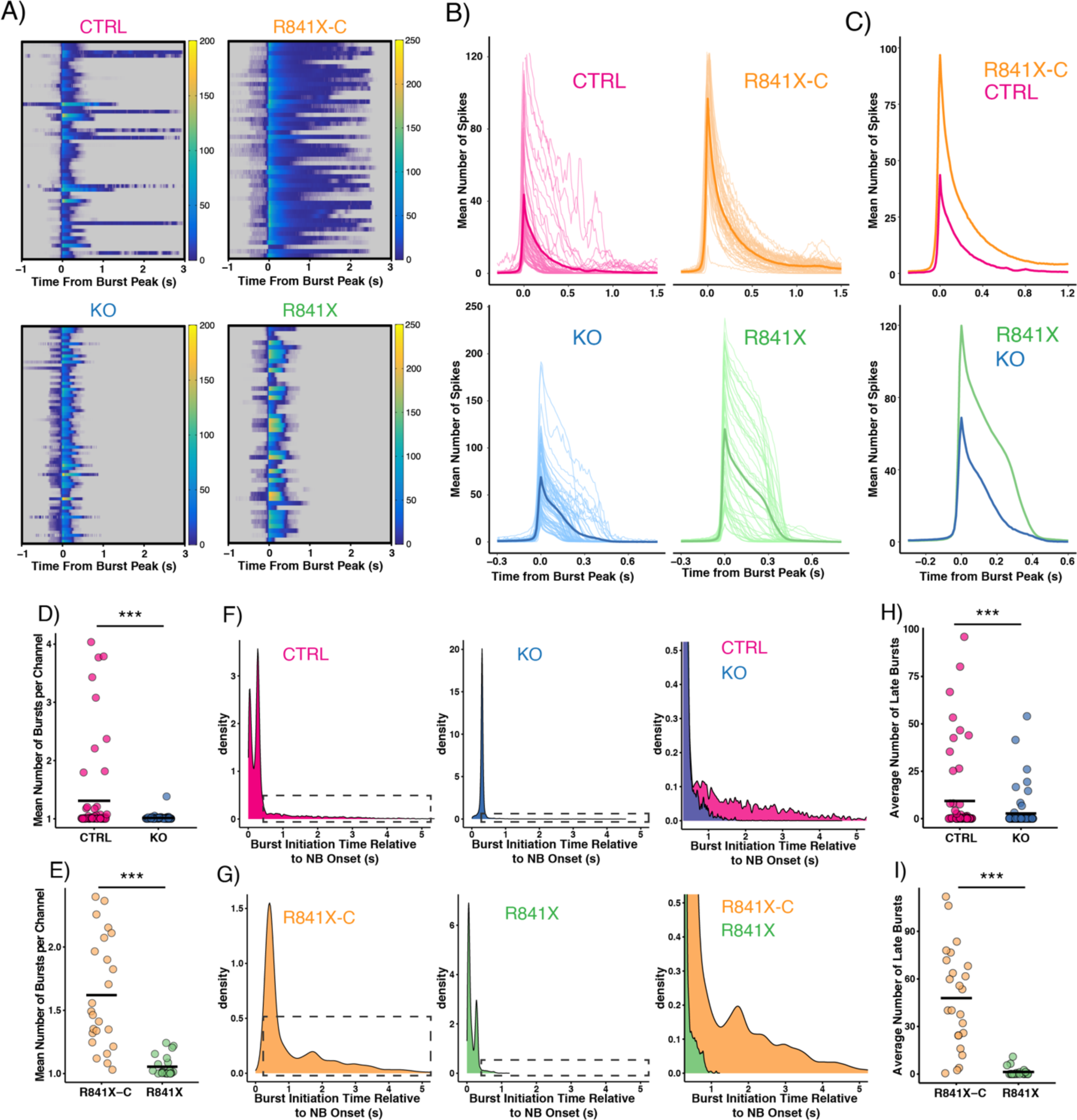
Intra-network burst firing dynamics are altered in *SHANK2* networks. (**A**) Heatmaps showing the average number of spikes per 10 ms bin within network bursts across all recordings for each cell line. Bins containing zero spikes are shown in grey to better distinguish them from periods of low-frequency spiking. (**B**) Line plots showing the average number of spikes per bin within network bursts for each cell line. Thin, light coloured lines show the average spikes per bin within each recording. The darker bolded line shows the average number of spikes per bin taken across all recordings. (**C**) Comparison of average network burst shapes (bolded lines in panel B) in control (top) and *SHANK2* (bottom) networks. (**D-E**) Mean number of bursts per bursting channel. (**F**) Smoothed gaussian kernel density estimates showing the distribution of single-channel burst initiation times relative to the time of network burst (NB) onset in *SHANK2* KO and CTRL and (**G**) *SHANK2* R841X and R841X-C networks. Right-most panel shows close up of boxed regions in top and middle plots, highlighting bursts which initiate more than 500 ms after network burst onset (i.e. late bursts). (**H-I**) Quantification of the percentage of late bursts per bursting channel. All spikes and bursts occurring outside of network burst events were omitted for calculations. Network recordings were taken from 6 independent differentiations for each cell line. ***P < 0.005; Mann-Whitney U test.

Interestingly, despite notable differences in the average network burst frequency, duration, and total spike content observed between the two control lines, their average network burst shapes were remarkably similar (Figure 4C, top), suggesting similar intra-network burst firing dynamics. Qualitatively, control network bursts appeared to consist of two distinct phases – a sharp increase in firing rate from network burst initiation to network burst peak, followed by a smooth exponential-like decay of the network-wide firing rate until network burst termination. Moreover, we observed that the network burst shapes of the two *SHANK2* networks were also qualitatively similar to one another, and appeared to differ from those of isogenic control networks in a consistent manner (Figure 4C, bottom). While the initial decay in *SHANK2* network firing rate following the burst peak appears to follow a similar trajectory to that of controls, this initial rapid decline in firing rate quickly transitions to a slower and more linear rate of decay. After this phase of extended high frequency firing, the rate of decay changes again, and the network-wide firing rate rapidly declines until burst termination.

We hypothesized that the characteristic *SHANK2* network burst shape could reflect a model where the prolonged high frequency firing in the initial phase of decay rapidly depletes neurons of available resources, leading to the subsequent crash in network-wide spiking activity and termination of the network burst (Huang et al., 2017). In contrast, the rapid exponential-like decline in control network firing rates seen immediately after the burst peak may provide individual cells within the network with sufficient time to replenish resources and initiate additional rounds of individual bursts to prolong the overall network burst event.

One prediction of this model, where individual control neurons have a better opportunity to replenish synaptic resources and burst again within the same extended network burst, is that they should have a greater number of bursts per single-channel recording than neurons in *SHANK2* networks. Consistent with this, we found that *SHANK2* KO and *SHANK2* R841X network bursts contained an average of 1.0 bursts per channel, while control and R841X-C networks contained a modest but significantly greater average of 1.6 (Figure 4D-E). Additionally, plotting the distribution of single-channel burst initiation times within network burst events revealed a greater density of bursts starting at later timepoints in control networks (Figure 4F-G). To quantify this, we classified single-channel bursts into two groups: “early bursts” (single-channel bursts which initiate within 500 ms of network burst onset) and “late bursts” (single-channel bursts which initiate greater than 500 ms after network burst onset). Both *SHANK2* KO and *SHANK2* R841X lines were found to have significantly fewer late bursts within network burst events than their respective isogenic controls (Figure 4H-I). Finally, we compared the distribution of interspike intervals (ISIs) within network bursts for *SHANK2* and control networks and found that *SHANK2* distributions were significantly more skewed towards shorter ISI values, indicative of higher intra-burst firing rate (P < 0.0001 for CTRL vs *SHANK2* KO distribution, P < 0.0001 for R841X-C vs *SHANK2* R841X distributions; One tailed, two-sample Kolmogorov-Smirnov test. Supplemental Figure 3).

Together, these results suggest that the majority of *SHANK2* network burst events primarily consist of a single, strong bursting event which persists for most of the network burst duration and encompasses the majority of network burst spikes. In contrast, control network bursts are less tightly structured and contain a greater diversity of single-channel bursts interspersed with low-frequency spiking.

### Machine learning classification of SHANK2 networks based on MEA metrics

Machine learning-based methods, such as linear classifiers, can be used as an unbiased approach to identify phenotypes in electrophysiological data (Hornauer et al., 2024). These methods are especially valuable when more than one electrophysiological feature carries information about the phenotype, and to be effective must be able to accurately distinguish between the *SHANK2* networks and the isogenic controls. By ranking the contributions of individual features, it should be possible to determine whether our supervised network analyses conducted with optimized network burst detection algorithms identify the most important features embedded in the activity metrics that can be read out from the automated MEA data analysis software provided by Axion systems.

We used the AceDimer machine learning classifier tool (see methods) which employs principles similar to Permutation Feature Importance (Altmann et al., 2010, Kaneko, 2022) and SHAP values (Lundberg and Lee, 2017). We provided it with a training set of raw Axion system metrics related to burst duration, synchrony, and network burst frequency and used this as a tool to identify the *SHANK2* and control networks in an independent dataset. The AceDimer method reached high performance levels with 85-90% accuracy at separating *SHANK2* R841X from the R841X-C isogenic controls, as well as at separating *SHANK2* KO networks and CTRL (Figure 5A, chance performance was 50%). To obtain an estimate of which features each classifier used to discriminate between the *SHANK2* and control networks, we ranked the features that both classifiers considered most informative (Figure 5B) by their contributions as percentages (Fig 5C). The top 6 feature contributions for the *SHANK2* R841X comparison related to network burst duration, synchrony (Area under Cross Correlation), and network burst frequency. This analysis provides additional support for our findings of network burst differences between *SHANK2* and controls, and suggests that classifiers may be a useful tool to distinguish between different network functional phenotypes.

**Figure 5.**
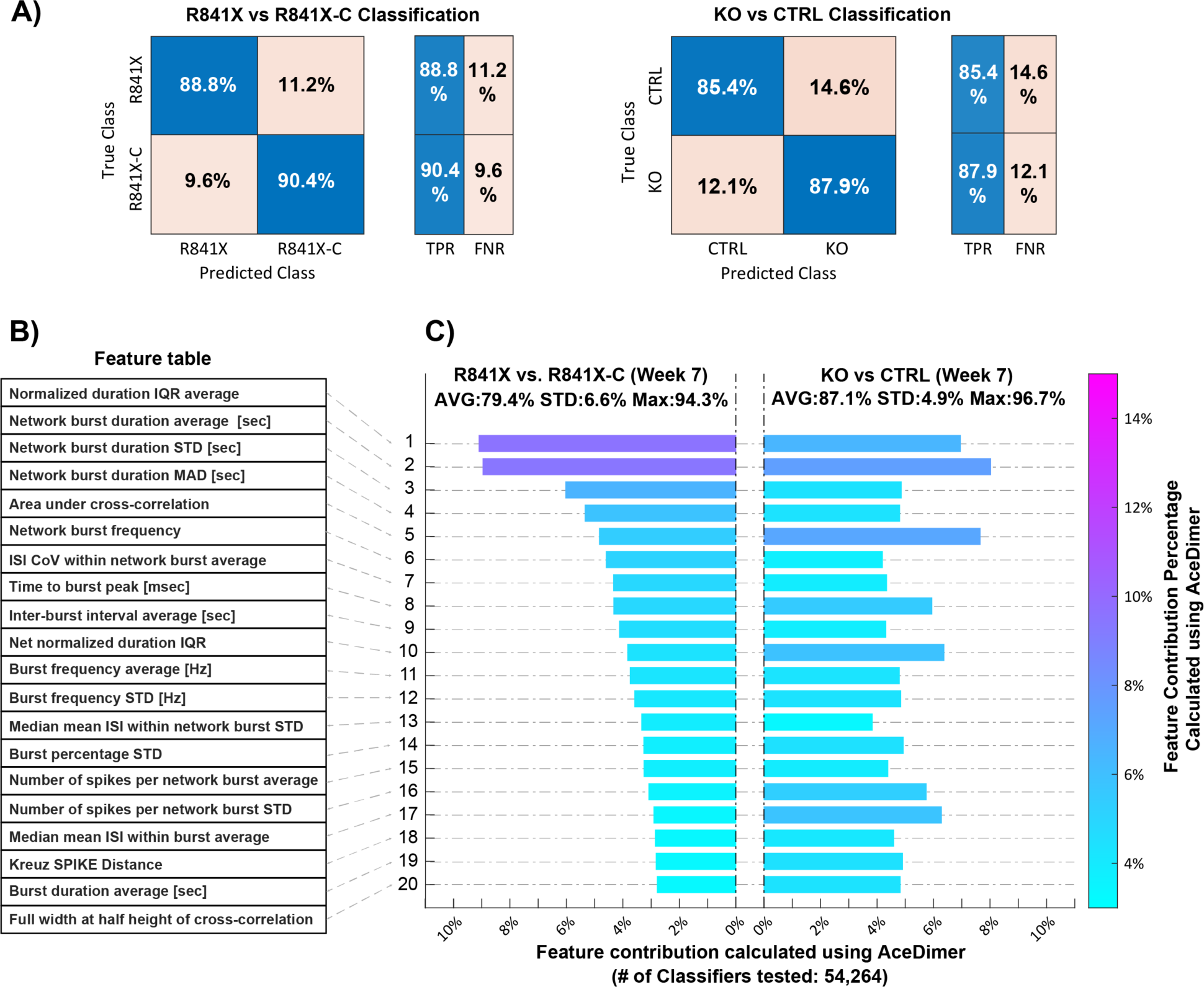
Machine learning classification of *SHANK2* network features. (**A**) Comparative Confusion Matrices for Classification Analyses of *SHANK2* R841X vs. R841X-C (top) networks and *SHANK2* KO vs. CTRL networks (bottom). Each matrix displays the predictive performance for each class in terms of accuracy percentages. True Positive Result (TPR), False Negative Result (FNR). (**B-C**) Network feature contributions to classification. Comparative Feature Contribution Analysis between *SHANK2* and control network features visualizes the normalized feature contribution percentages for twenty key features across the two study groups. Contributions are calculated using the AceDimer algorithm, demonstrating the influence of each feature within the classification model. Each bar represents the feature contribution percentage to classification accuracy, with colour intensity indicating relative importance.

### Calcium-dependent reverberating super bursts in control and SHANK2 networks

RSBs are network events consisting of a long initial high amplitude burst followed by high frequency smaller amplitude minibursts. Such events are found in control networks and they are particularly frequent in Rett syndrome networks with mutations of the *MECP2* gene (Pradeepan et al., 2024). Here we explore whether RSBs also occur in *SHANK2* cultures. RSBs were detected in CTRL and R841X-C cultures beginning at week 4 and week 5 of development, respectively, and persisted until the end of our recording time course at week 8. RSBs were very rarely detected in *SHANK2* R841X wells at week 7 and week 8 (n=1 and n=2, respectively), but were never detected in *SHANK2* KO networks (Figure 6A-C).

**Figure 6.**
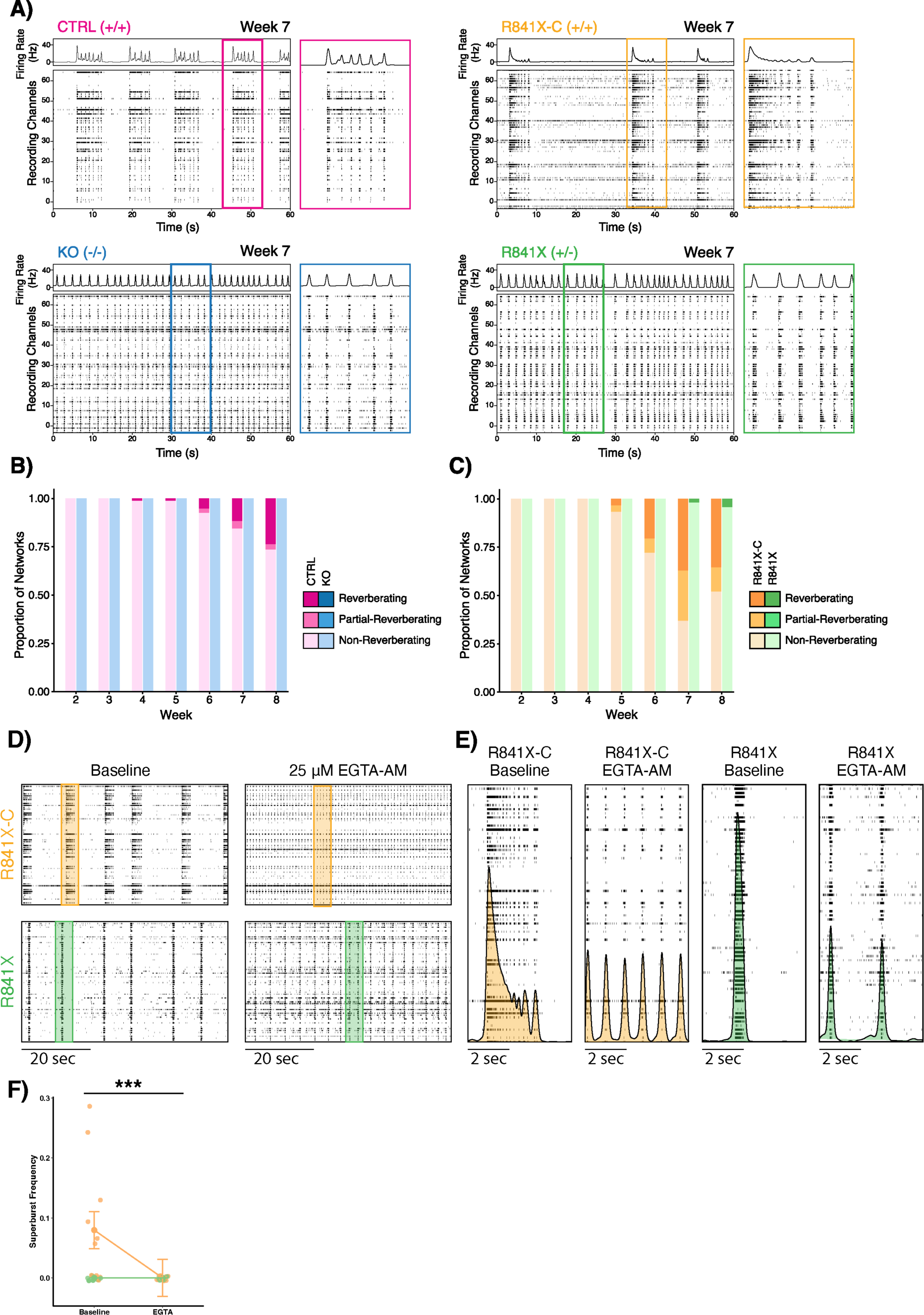
Detection and quantification of Ca^2+^ sensitive reverberating super bursts in control networks. (**A**) Representative raster plots from *SHANK2* and control network recordings showing the appearance of RSBs in control (top) but not *SHANK2* (bottom) networks. (**B**) Quantification of the proportion of reverberating, partial-reverberating, and non-reverberating networks from week 2 – week 8 of development in *SHANK2* KO and CTRL cultures, and in (**C**) *SHANK2* R841X and R841X-C cultures. (**D**) Representative raster plots for *SHANK2* R841X and control R841X-C networks before and after treatment with 25 µM EGTA-AM. (**E**) Closeups of shaded regions in (D) showing network burst structure. (**F**) Quantification of reverberating super burst frequency in networks before and after treatment with EGTA-AM. ***P < 0.005; Mann-Whitney U test.

We previously showed that RSBs occurring in networks of *MECP2* null neurons were abolished by treatment with the membrane-permeable Ca^2+^ chelator ethylene glycol tetraacetic acid acetoxymethyl ester (EGTA-AM) (Pradeepan et al., 2024). A different study performed on organoids demonstrated GABAergic dependence of nested oscillations, a phenomenon similar to RSBs, using bicuculline (Trujillo et al., 2019). Accordingly, we next treated *SHANK2* R841X and control R841X-C networks with either 10 µM bicuculline or 25 µM EGTA-AM to investigate what effect this had on the presence of RSBs. Bicuculine treatment did not abolish RSBs in reverberating R841X-C networks (Supplemental Figure 4A-B). However, bicuculline treatment did significantly increase the frequency of RSBs in R841X-C cultures and significantly decreased their duration (Supplemental Figure 4C). These bicuculine side effects were seen previously in treated control and *MECP2* null networks (Pradeepan et al., 2024) and may be due to off target effects on potassium channels (Khawaled et al., 1999). In contrast, treatment with EGTA-AM completely abolished RSBs in reverberating R841X-C networks (Figure 6D-E) corroborating the previous results that RSBs are indeed dependent on intracellular Ca^2+^ (Pradeepan et al., 2024).

### Chronic mGluR stimulation with DHPG rescues hypersynchronous SHANK2 networks

We previously showed that chronic treatment of *SHANK2* R841X neurons with 10 µM of the group 1 mGluR agonist DHPG was able to rescue the observed increased dendrite length phenotype (Zaslavsky et al., 2019). We next asked if chronic DHPG treatment could similarly rescue or improve any aspects of the *SHANK2* electrophysiological network phenotype. We treated *SHANK2* R841X and isogenic R841X-C networks by adding 10 µM DHPG with every culture media change from week 1 – week 8 and recorded spontaneous network activity. Chronic DHPG treatment did not significantly impair network formation as robust spiking activity and network burst formation was observed up to week 8 of development in DHPG-treated networks (Figure 7A, Supplemental Figure 5A). In addition, DHPG treatment had no impact on MFR in either *SHANK2* R841X or control R841X-C lines (Supplemental Figure 5B).

**Figure 7.**
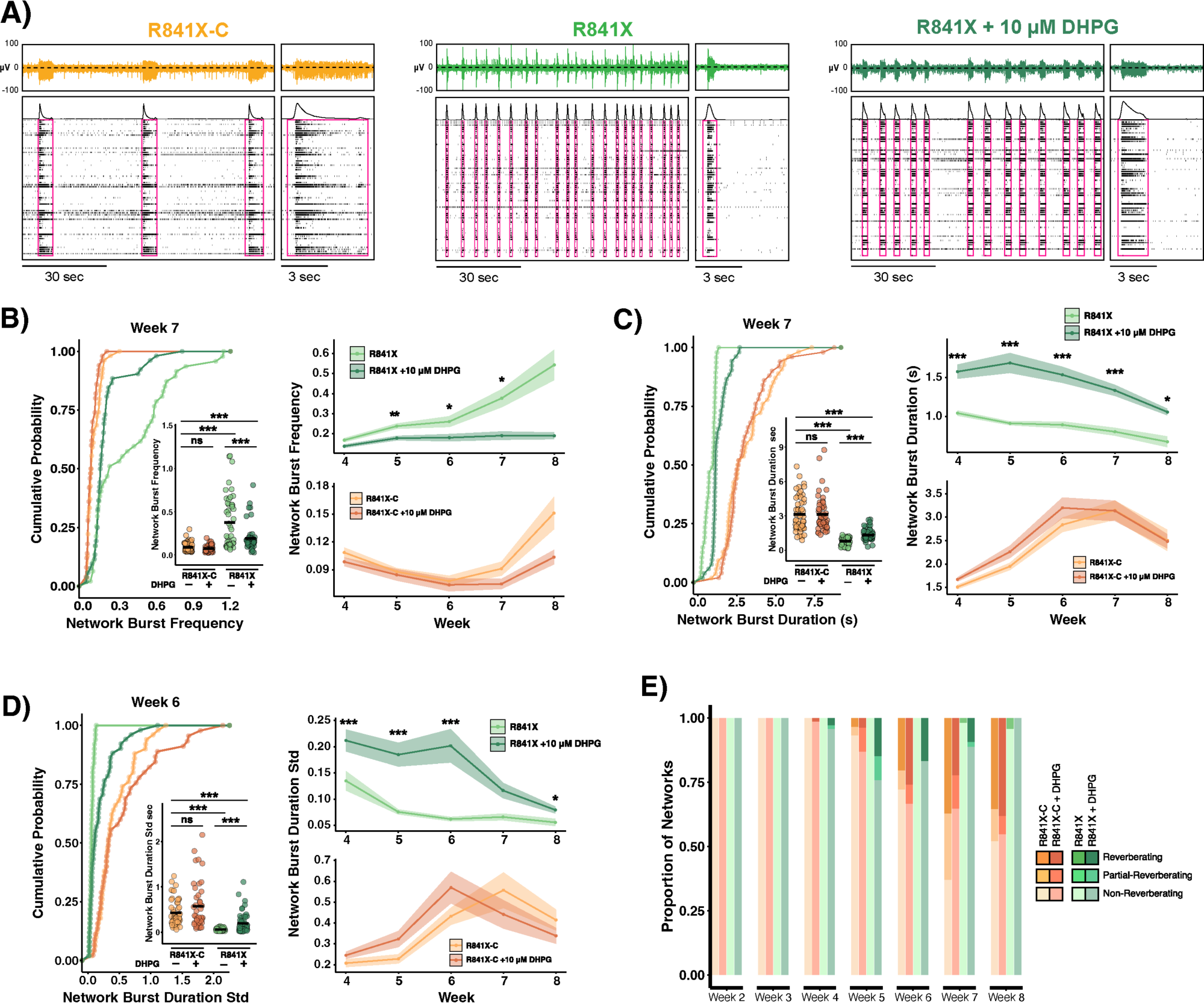
DHPG improves *SHANK2* network burst frequency, duration and RSB detection. **(A)** Representative raster plots and extracellular voltage traces from isogenic *SHANK2* R841X and control R841X-C cultures with and without DHPG treatment. (**B**) Quantification of network burst frequency, (**C**) network burst duration and (**D**) the standard deviation of network burst durations in *SHANK2* R841X and control R841X-C networks, untreated or treated with 10 µM DHPG. (**E**) Quantification of the proportion of reverberating, partial-reverberating, and non-reverberating networks from week 2 – week 8 of development in untreated and DHPG-treated *SHANK2* R841X and R841X-C networks. Network recordings were taken from 6 independent differentiations for each cell line. *P < 0.05, **P< 0.01, ***P < 0.005; ns, not significant; Mann-Whitney U test with Benjamini-Hochberg correction for multiple testing.

*SHANK2* R841X networks treated with DHPG exhibited a significant reduction in network burst frequency, and a significant increase in network burst duration when compared to untreated *SHANK2* R841X networks at all time points investigated (Figure 7B-C). Network burst durations in *SHANK2* networks also became more variable with DHPG treatment, particularly at early time points (Figure 7D). This response was specific to *SHANK2* networks, as DHPG treatment had no impact on network burst frequency, duration, or duration variability in control R841X-C networks at all time points investigated (Figure 7B-D). Interestingly, DHPG treatment also appeared to increase the proportion of *SHANK2* R841X wells displaying patterns of RSB activity (Figure 7E). However, while persistent exposure to 10 µM DHPG in the culture media was able to improve these aspects of the *SHANK2* network phenotype, it did not fully restore these metrics to the level of untreated control networks.

Next, we turned our attention to intra-network burst firing dynamics. We again plotted the average network burst shape for each recording and found that *SHANK2* R841X networks appeared to have a variable response to DHPG treatment. While a subset of wells retained the characteristic *SHANK2* network burst shape with a slower decline in firing rates after the burst peak, the remaining networks displayed network burst shapes that more closely resembled controls with firing rates that decayed exponentially following the burst peak (Figure 8A-C). Examining the distribution of single-channel burst start times within network burst events, *SHANK2* R841X networks treated with DHPG now displayed a prominent second peak of burst start times initiating 1 – 2 seconds after network burst onset, similar to the R841X-C controls (Figure 8D). This was quantified as a significant 18.5 fold increase in the percentage of single-channel bursts occurring more than 500 ms after network burst onset (i.e. “late bursts”), from a mean of 2% in untreated *SHANK2* R841X networks to a mean of 37% in DHPG treated networks (Figure 8E). Moreover, we found a significant 1.5-fold increase in the number of bursts per bursting channel (untreated mean = 1.0, treated mean = 1.5; Figure 7F). DHPG treatment was also found to reduce the number of network burst spikes occurring within single-channel bursts (Figure 8G) and increased the average within-burst ISI (Figure 8H).

**Figure 8.**
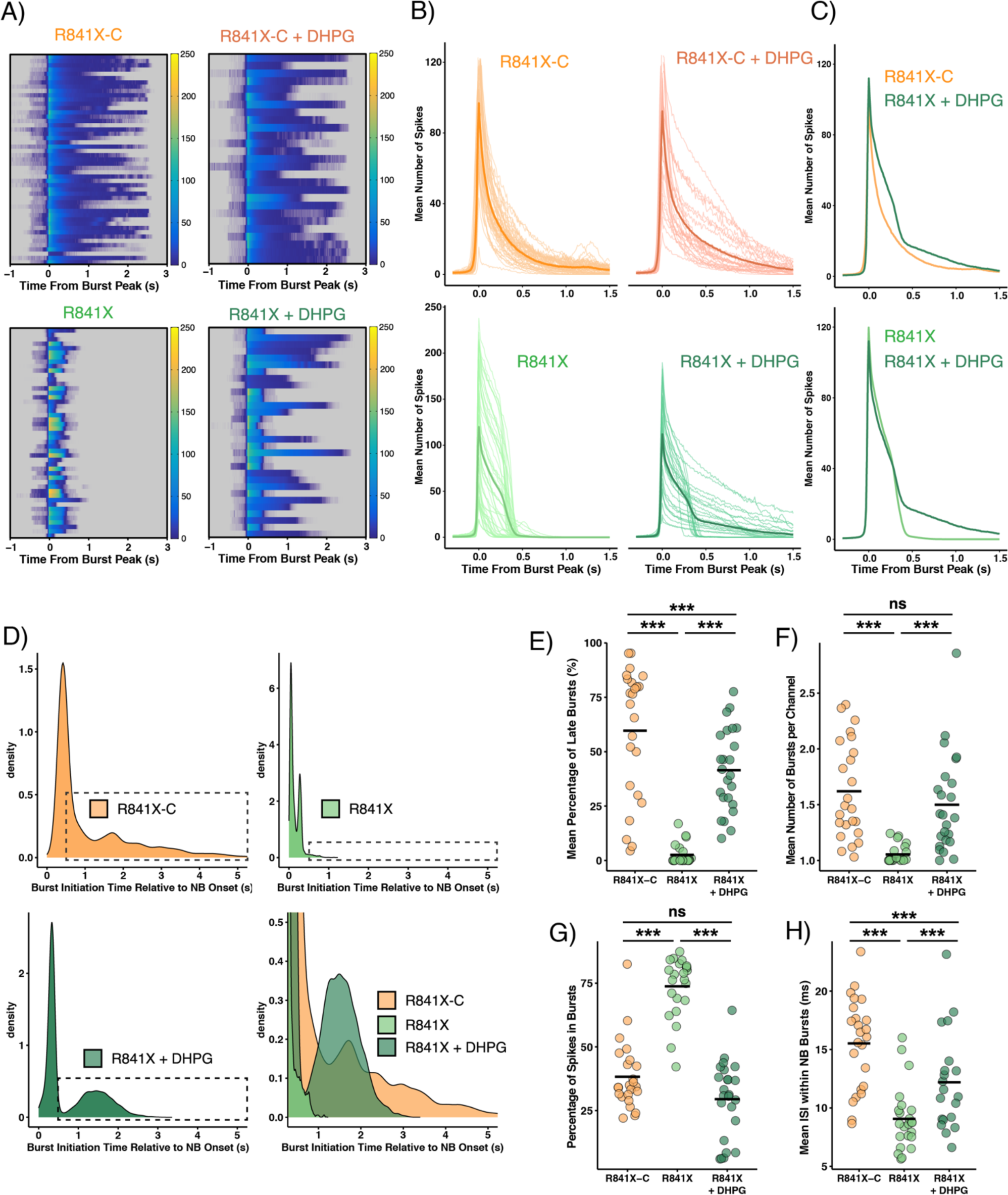
DHPG improves intra-network burst firing dynamics in *SHANK2* R841X networks. (**A**) Heatmaps showing the average number of spikes per 10 ms bin within network bursts across all recordings for each cell line. Bins containing zero spikes are shown in grey to better distinguish them from periods of low-frequency spiking. (**B**) Line plots showing the average number of spikes per bin within network bursts for each cell line. Thin, light coloured lines show the average spikes per bin within each recording. The darker bolded line shows the average number of spikes per bin taken across all recordings. (**C**) Comparison of average network burst shapes (bolded lines in panel B) between untreated control R841X-C and DHPG treated *SHANK2* R841X networks (top), and between untreated and DHPG treated *SHANK2* R841X networks (bottom). (**D**) Smoothed gaussian kernel density estimates showing the distribution of single-channel burst initiation times relative to the time of network burst (NB) onset. Bottom-right panel shows close up of boxed regions in top and bottom left plots, highlighting bursts which initiate more than 500 ms after network burst onset (i.e. late bursts). (**E**) Quantification of the percentage of late bursts per bursting channel. (**F**) Mean number of bursts per bursting channel. (**G**) Percentage of spikes in bursts. (**H**) Mean ISI of single-channel bursts that occur within network bursts. All spikes and bursts occurring outside of network burst events were omitted for calculations in (E-H). Network recordings were taken from 6 independent differentiations for each cell line. ***P < 0.005; ns, not significant; Mann-Whitney U test.

Finally, we assessed if DHPG treatment could rescue the increased synchrony displayed by *SHANK2* networks. Representative rasters and CorSE output are shown in Figure 9A-B. CorSE analysis of network synchronization revealed a significant shift in the distribution of synchrony scores in DHPG-treated *SHANK2* R841X networks. Notably, the prominent secondary peak centered around CorSE values of approximately 0.6 seen in untreated *SHANK2* R841X networks was noticeably flattened by DHPG treatment (Figure 9C-D). While DHPG-treated *SHANK2* R841X networks were found to have a slight increase in the number of correlations with CorSE scores in the range of 0.8 – 1.0 (Figure 9D), the overall distribution of correlation strengths in treated networks was significantly shifted towards lower synchrony values (P < 0.0001, one tailed, two-sample Kolmogorov-Smirnov test). This shift could further be quantified as a significant 1.9-fold reduction in the average mean CorSE synchrony score in DHPG-treated *SHANK2* R841X networks (Figure 9E). Importantly, DHPG-treated *SHANK2* R841X networks were restored to the level of the isogenic controls as they showed no significant difference from the mean synchrony scores of R841X-C cultures. Additionally, DHPG treatment resulted in a significant 3.3-fold reduction in the number of strong correlations (CorSE score > 0.5) in *SHANK2* R841X networks (Figure 9F). Again, DHPG-treated *SHANK2* R841X networks were equivalent to the isogenic controls, with no significant difference detected in the number of strong correlations from R841X-C cultures. Taken together, these results show that chronic group 1 mGluR agonism with 10 µM DHPG was successful in rescuing the hypersynchronous phenotype displayed by *SHANK2* R841X networks.

**Figure 9.**
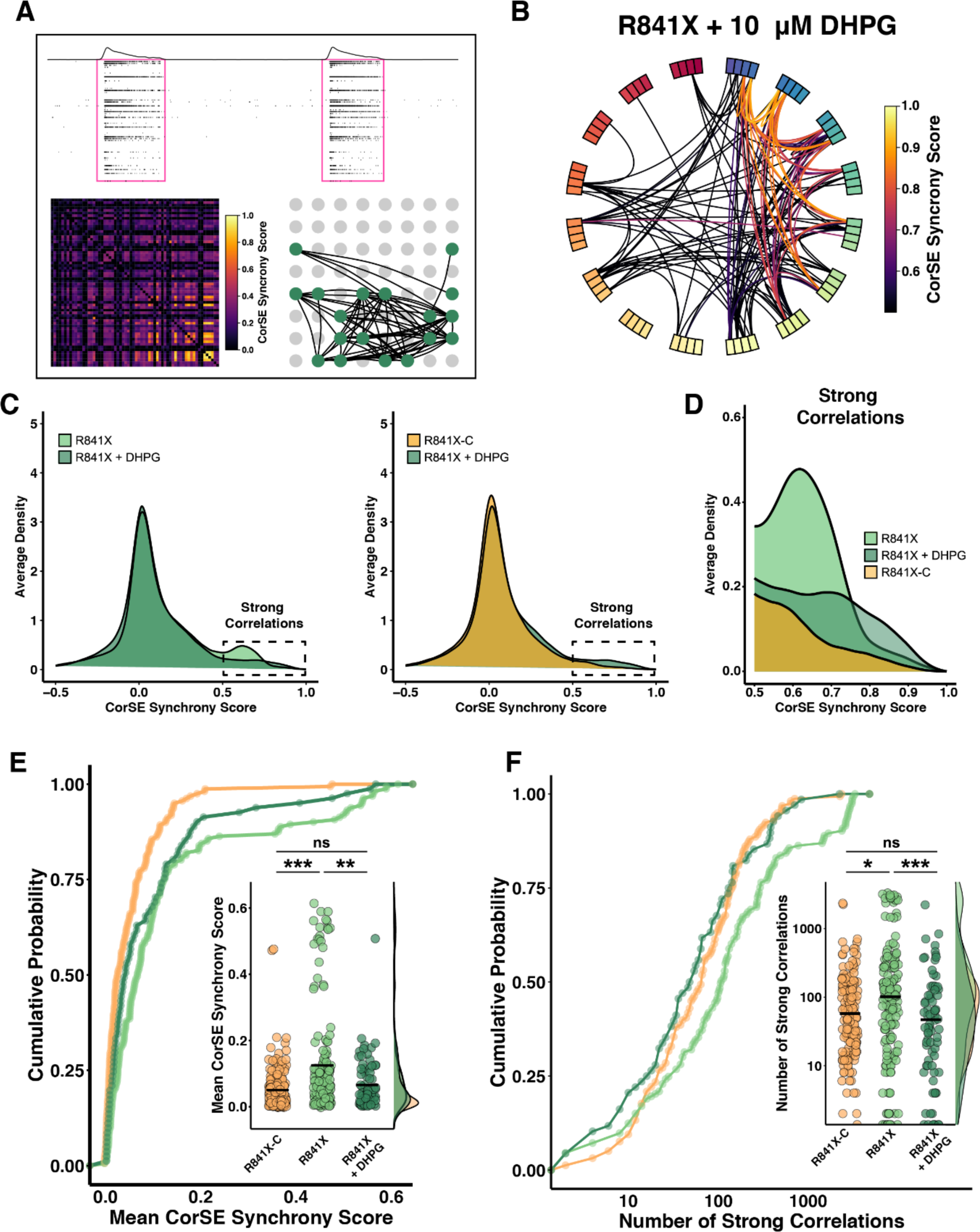
DHPG rescues mean CorSE synchrony score and distribution of correlation strengths in *SHANK2* R841X networks. (**A**) Representative CorSE output for DHPG-treated *SHANK2* R841X networks. Sub-panels show a raster plot of 15 seconds activity from the 5-minute recordings and corresponding correlation matrix showing the synchrony score for each electrode pair (bottom left). The bottom right panel shows the 8 x8 electrode grid with an overlaid a connectivity map for the recording. Each edge represents a connection between two electrodes with a synchrony score > 0.5. (**B**) Circular connectivity plots from representative recordings showing all pairwise correlations with a strength > 0.5. Boxes around the circumference of the circle represent electrodes and edges represent connections. Edge colour indicates the strength of connection. (**C**) Smoothed gaussian kernel density estimates comparing the distribution of correlation strengths across all recordings at week 7 and 8. (**D**) Close up of boxed regions marked in (C), highlighting strong correlations (CorSE score > 0.5). (**E**) Quantification of mean CorSE synchrony score and (**F**) the number of strong correlations at week 7. *P < 0.05, **P< 0.01, ***P < 0.005; ns, not significant; Mann-Whitney U test.

## DISCUSSION

In this study, we provided a thorough phenotypic characterization of *SHANK2* neuronal network activity *in vitro*. We found that *SHANK2* networks were hyperactive with increased MFR over controls. Moreover, *SHANK2* networks were hypersynchronous with increased mean CorSE synchrony scores and an increased number of strong pairwise correlations between recording channels. At the network activity level, these traits manifested as an increased frequency of network bursts, which were shorter and more regular in duration than controls. Examination of intra-network burst firing dynamics revealed a characteristic *SHANK2* network burst shape which was consistent across *SHANK2* R841X and *SHANK2* KO cultures. This network burst shape was characterized by a significantly slower decline in network-wide firing rates after the burst peak, followed by a rapid termination of network activity. We found that this characteristic network burst shape, at least in part, reflected a decreased ability of *SHANK2* networks to drive additional rounds of single-channel burst firing following the initiating burst of activity. Consistent with this, Ca^2+^ dependent RSBs rarely occurred in *SHANK2* networks but were present in controls. Finally, we showed that treatment of *SHANK2* R841X networks with DHPG was able to fully rescue the hypersynchronous *SHANK2* network phenotype.

### Intra-network burst dynamics and network burst shape

Our finding that *SHANK2* and control neurons had characteristic network burst shapes is intriguing, as this appears to be robust to substantial within-genotype differences in the gross firing patterns displayed between lines when examined at larger timescales. Indeed, the network burst shapes displayed by individual cell lines was remarkably consistent across multiple neuronal differentiations and platings, and were observed to be robust to significant changes in the overall amplitude of intra-network burst firing rates. Characterization of intra-network burst dynamics could thus prove to be a useful approach for disease phenotyping studies, where batch effects and plate-to-plate variability in firing metrics are a known challenge for the field (Mossink et al., 2021).

We hypothesized that the characteristic network burst shapes produced by *SHANK2* networks could reflect a model consistent with synaptic fatigue, where the intense firing within *SHANK2* network events rapidly depletes the energy or synaptic resources in single neurons required to sustain the prolonged elevated firing rate across an excitatory-only network. This leads to a sudden and rapid decrease in firing rate immediately preceding network burst termination as synaptic resources are depleted. The fact that DHPG rescued many aspects of the hyperconnected phenotype suggests that *SHANK2* variants affect the synaptic machinery that allows fast electrochemical transmission during network bursts. Moreover, our finding that DHPG treatment concurrently restores wild-type-like network burst dynamics in a subset of *SHANK2* networks provides some supporting evidence that these characteristic bursting dynamics are influenced by coupling strength within the network.

It should be noted that differences in the intrinsic properties of single neurons could also influence the network burst shapes (e.g., spike frequency adaptation) observed in our excitatory-only networks. Future studies using *in silico* network models could be used to dissect the contributions of intrinsic and synaptic properties to network bursting behaviour, and to test the predictions we have made here.

### Network reverberating super bursts

Our finding that RSB frequency is decreased in *SHANK2* networks demonstrates the potential utility of RSB quantification in disease phenotyping assays. The presence of RSBs in network recordings was also found to segregate by genotype in *MECP2* null Rett syndrome models (Pradeepan et al., 2024). Interestingly, RSBs were found more frequently in *MECP2* null networks than controls, while they were rarely detected in *SHANK2* networks. The emergence of RSBs in networks prone to generating them appears to follow a non-random trajectory during network development, with RSBs increasing in frequency over development. Therefore, bursting patterns dominated by RSBs might thus reflect a particular stage of network development. Based on all this, RSBs could be useful indicators of the network developmental trajectory.

In studies of primary mouse neurons, this reverberation stage has been noted to be transitory in nature, with patterns of long super bursts transitioning to more rapid, short duration burst firing later in network development (Wagenaar et al., 2006). One possibility is that the ability of excitatory-only networks to generate RSBs may be tied to network connectivity levels, which change as networks mature or as a function of gene variants. At the early stages of network development, networks are marked by nascent synapses, unable to produce successive rounds of RSBs. As networks develop, synaptic strength increases, promoting noisy asynchronous neurotransmitter release following an initial burst of activity, which emerges as RSBs. RSBs, with their sustained activation of postsynaptic receptors, may be critical in synaptic plasticity of the network. Over time, as the strength of synapses increases, the initiation burst becomes more rapid and strong, depleting neurons of available neurotransmitter resources to a degree where additional rounds of rapid burst firing cannot be supported without a sufficiently long refractory period. Therefore, if RSBs are taken as a reflection of underlying connectivity strengths within a network rather than a pathological state, the discordant findings between our previously reported *MECP2* null networks and *SHANK2* networks are less conflicting. In *MECP2* null networks, which are hypoconnected, networks may persist longer in a firing regime dominated by RSBs. On the other hand, *SHANK2* networks, which are hyperconnected and hypersynchronous, might bypass the RSB stage or transit through it at a rate which makes it difficult to detect on a biweekly recording schedule. In support of this, in our present study, the cell line which was most prone to generating RSBs (R841X-C) also had the lowest mean CorSE synchrony scores of the four lines we investigated, indicating lower levels of functional connectivity within these networks. Moreover, the lower levels of functional connectivity seen in *SHANK2* R841X networks treated with DHPG also coincided with an increase in the presence of RSBs observed in these networks.

### DHPG rescues hypersynchronous SHANK2 networks

DHPG treatment was most effective in rescuing differences in *SHANK2* network synchrony. Activation of mGluRs is associated with the induction of long-term depression and homeostatic synaptic scaling. Rescues of hyperconnectivity in *SHANK2* networks by DHPG could be reflecting an ability of the drug to successfully restore appropriate synaptic connectivity levels within the network. In support of this, we found that mean CorSE synchrony scores and the number of strong correlations were fully rescued by DHPG treatment, indicating a return to control levels of coupling strength. However, our finding that some activity metrics such as MFR were unaffected by DHPG treatment, while others such as network burst frequency and network burst duration saw partial rescue, would suggest that additional factors beyond average coupling strength contribute to the *SHANK2* network phenotype. Future experiments using *in silico* network models (Doorn et al., 2023) could prove useful in delineating the identities of these additional contributing factors.

## Acknowledgements

This study was funded by grants from Simons Foundation Autism Research Initiative (Grant No. 514918 to JE and JMT); the Ontario Brain Institute (POND Network to JE); John Evans Leaders Fund/Canada Foundation for Innovation (JELF/CFI) and Ontario Research Fund (to JE); Canada Research Chair (Tier 1) in Stem Cell Models of Childhood Disease (to JE); the Beta Sigma Phi International Endowment Fund (to JE); BrainsCAN at Western University through the Canada First Research Excellence Fund (to KSP and JMT). Trainee support was provided by a Natural Sciences and Engineering Research Council (NSERC) Doctoral Scholarship (to KSP); and an Autism Speaks Predoctoral Award, Ontario Graduate Student (OGS) Award, and David Stephen Cant Scholarship in Stem Cell Research (to FPM).

## Author Contributions

Conceptualization, FPM, KSP, JMT, JE; Methodology, FPM, KSP, MK, WW; Software, FPM, MK; Formal Analysis, FPM, KSP, MK, NM; Investigation, FPM, KSP, MK, WW, NM, AP; Writing – Original Draft, FPM, KSP, MK, WW, NM, JMT, JE; Writing – Review and Editing, FPM, KSP, MK, WW, NM, AP, JMT, JE; Visualization, FPM, KSP, MK; Supervision, JMT, JE; Funding Acquisition, FPM, KP, JMT, JE.

