## Supplemental Figures for "Hypersynchronous iPSC-derived *SHANK2* neuronal networks are rescued by mGluR5 agonism"

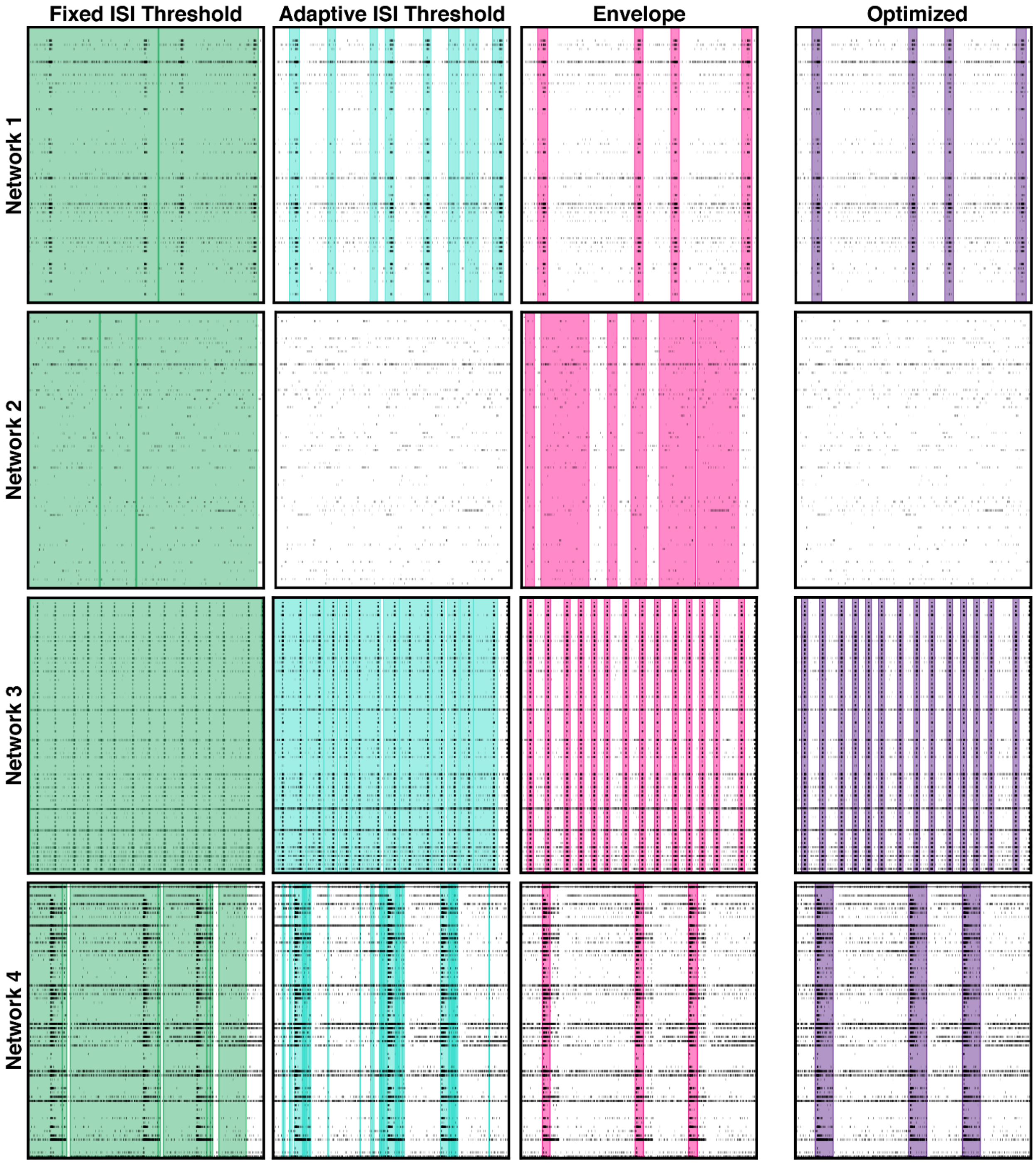


**Supplemental Figure 1. Common network burst detection settings can lead to spurious network burst calls.** Network burst calls from fixed ISI threshold (green), adaptive ISI threshold (teal), and envelope based (purple) network burst detection algorithms overlaid on raster plots from 4 representative network recordings. Optimized network burst detection for each network shown in purple. Shaded regions indicate network burst boundaries determined by each algorithm.

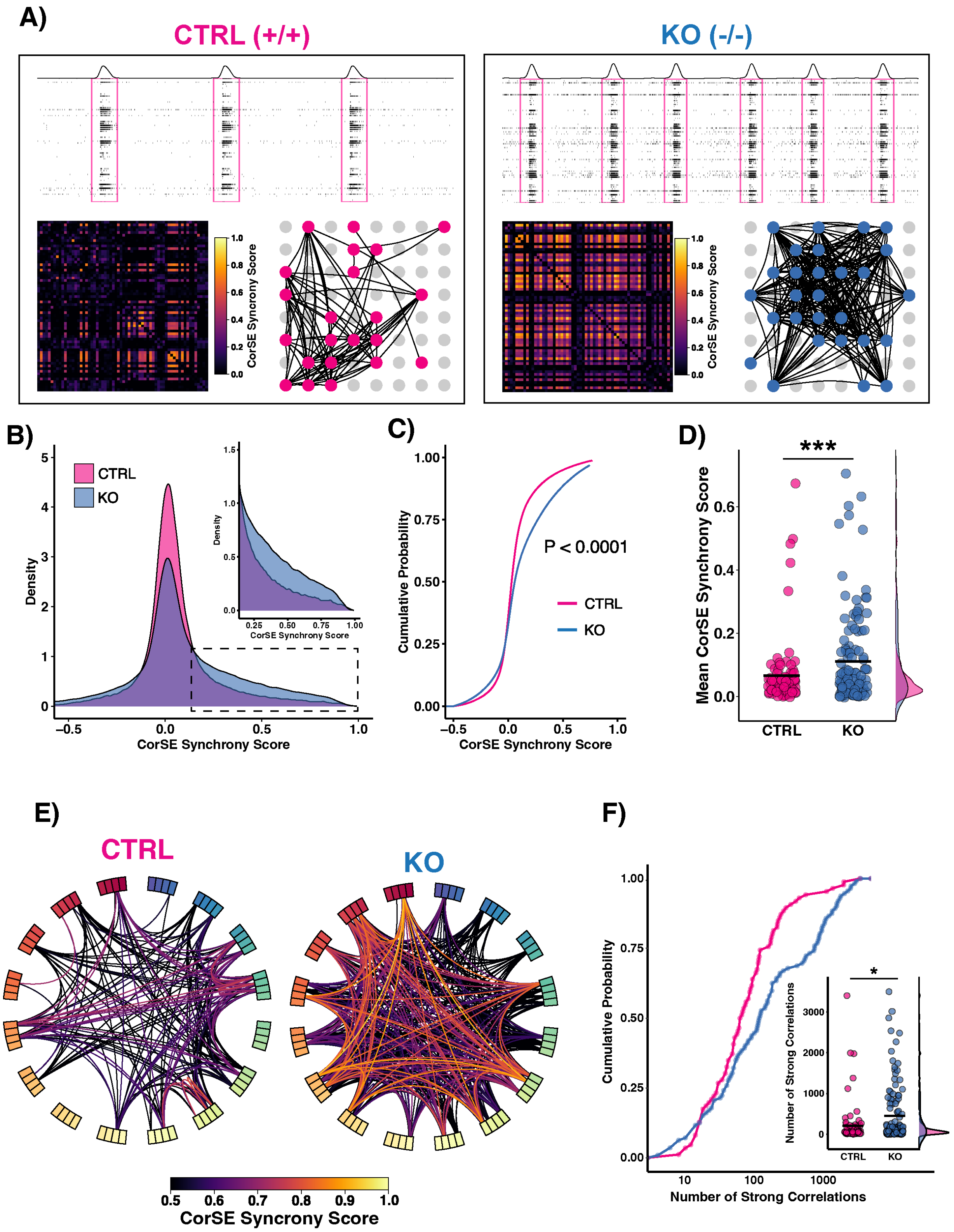

A)
CTRL (+/+)
KO (-/-)
CorSE Syncrony Score
1.0
0.8
0.6
0.4
0.2
0.0
B)
CTRL
KO
Density
CorSE Synchrony Score
C)
Cumulative Probability
P < 0.0001
CTRL
KO
CorSE Synchrony Score
D)
***
Mean CorSE Synchrony Score
CTRL
KO
E)
CTRL
KO
0.5 0.6 0.7 0.8 0.9 1.0
CorSE Syncrony Score
F)
Cumulative Probability
*
Number of Strong Correlations
CTRL
KO
10 100 1000
Number of Strong Correlations

**Supplemental Figure 2. Increased synchrony of *SHANK2* KO networks as measured by correlated spectral entropy.** (**A**) Representative CorSE output for *SHANK2* R841X and control R841X-C networks. Each sub-panel shows a raster plot of 15 seconds activity from the 5-minute recordings and corresponding correlation matrix showing the synchrony score for each electrode pair (bottom left). The bottom right panel shows the 8 x8 electrode grid with an overlaid a connectivity map for the recording. Each edge represents a connection between two electrodes with a synchrony score > 0.5. Coloured nodes indicate electrodes involved in these connections and are shown at their appropriate spatial location on the electrode grid. (**B**) Smoothed gaussian kernel density estimates comparing the distribution of connection strengths across all recordings at week 7 and 8 in *SHANK2* R841X and R841X-C networks. (**C**) Empirical cumulative distribution functions comparing CorSE synchrony scores in R841X-C and *SHANK2* R841X networks. (**D**) Quantification of mean synchrony scores in all recorded networks. (**E**) Circular connectivity plots from representative recordings showing all pairwise correlations with a strength > 0.5. Boxes around the circumference of the circle represent electrodes and edges represent connections. Edge colour indicates the strength of connection. (**F**) *SHANK2* networks show increased number of strong correlations than controls. n = 112 for CTRL, n = 131 for KO. Network recordings were taken from 6 independent differentiations for each cell line). *P < 0.05, **P< 0.01, ***P < 0.005; Anderson-Darling k-samples test.

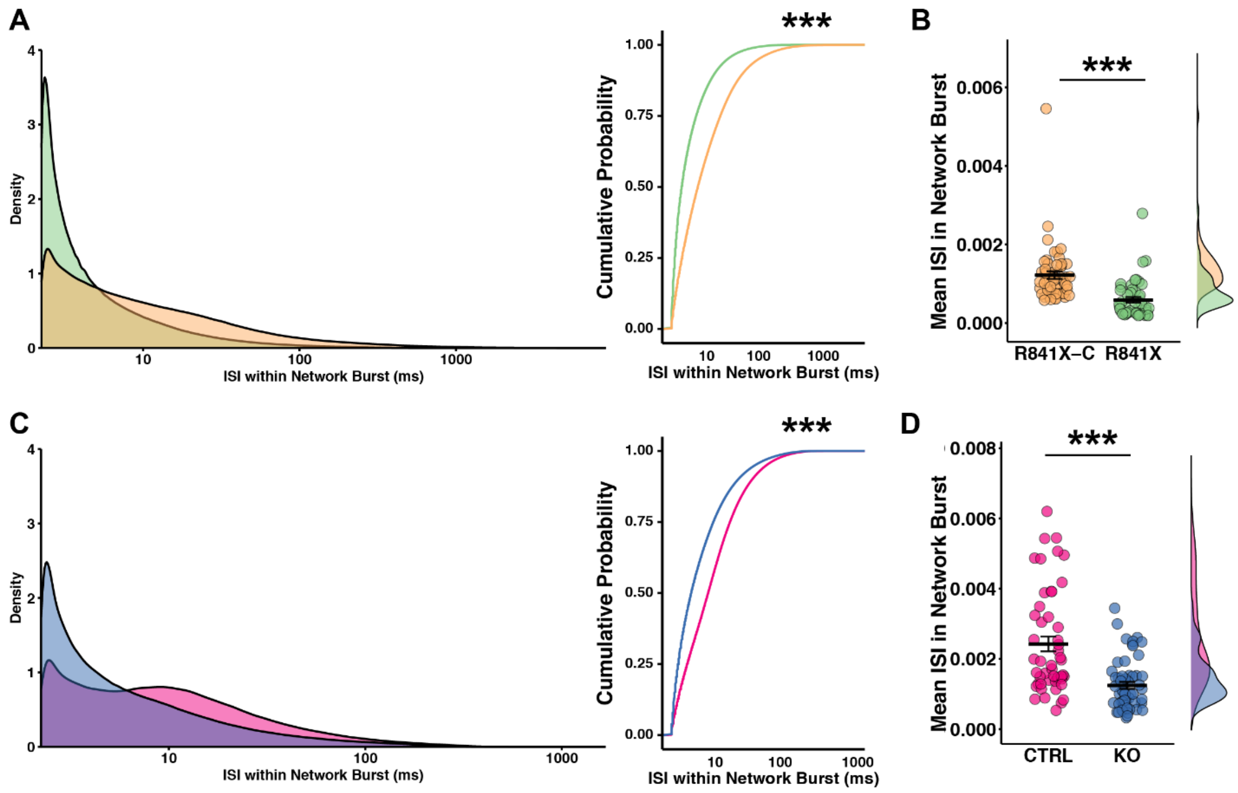


**Supplemental Figure 3. Distribution of ISIs in *SHANK2* network bursts are skewed towards shorter durations.** (**A,C**) Distribution of intra-network burst ISI values in *SHANK2* R841X (**A**) and *SHANK2* KO (**C**) show a significant skew towards shorter ISI values when compared to their respective isogenic controls. Smoothed gaussian kernel density estimates (left) and empirical cumulative distribution functions (right) of intra-network burst ISIs recorded at week 7 are shown. (**B,D**) *SHANK2* networks have lower mean network burst ISI values at week 7. Cossbars indicate the mean of each group and margin density plots show the distribution of datapoints (n = 54 for R841X-C, n = 47 for R841X, n = 60 for CTRL, n = 73 for KO. Network recordings were taken from 6 independent differentiations for each cell line). ***P < 0.005; one tailed, two-sample Kolmogorov-Smirnov test for (A) and (C). Mann-Whitney U test with Benjamini-Hochberg correction for multiple testing for (B) and (D).

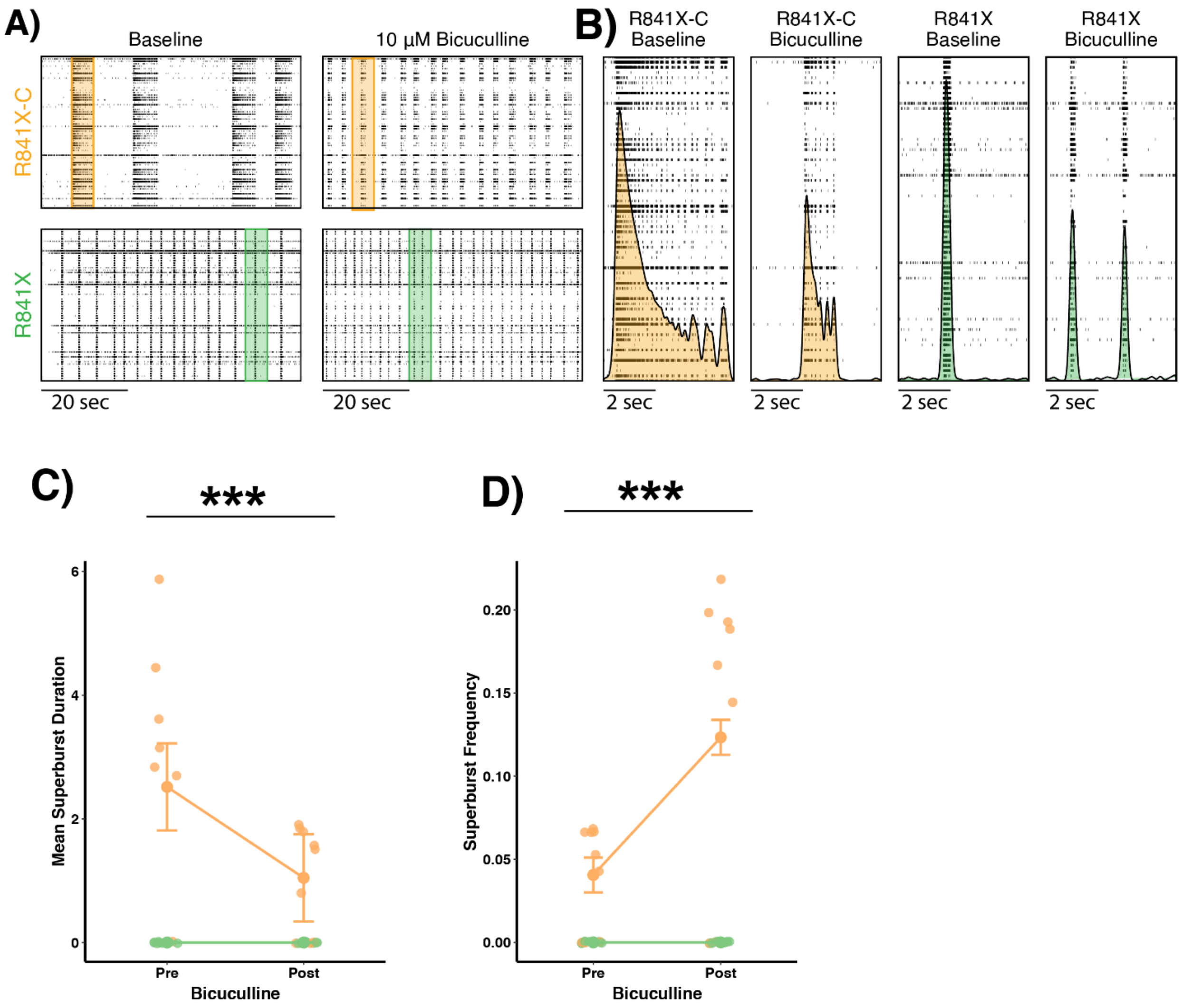


**Supplemental Figure 4. Bicuculline treatment does not eliminate RSBs in *SHANK2* R841X-C networks.** (**A**) Representative raster plots for *SHANK2* R841X and control R841X-C networks before and after treatment with 10 µM bicuculline. (**B**) Close-ups of shaded regions in (A) showing network burst structure. (**C**) Quantification of reverberating super burst frequency and (**D**) reverberating super burst duration in *SHANK2* R841X and R841X-C networks before and after treatment with bicuculline. ***P < 0.005; Mann-Whitney U test.

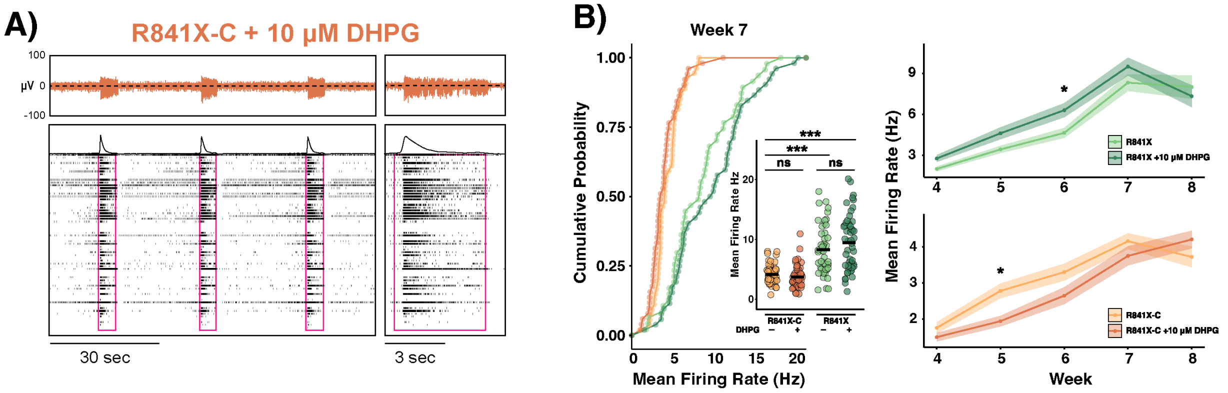


**Supplemental Figure 5. DHPG treatment has no impact on MFR in *SHANK2* networks.** (**A**) Representative raster plots and extracellular voltage traces from R841X-C cultures treated with DHPG. (**B**) Quantification of mean firing rate in treated and untreated cultures. Network recordings were taken from 6 independent differentiations for each cell line. *P < 0.05, **P< 0.01, ***P < 0.005; ns, not significant; Mann-Whitney U test with Benjamini-Hochberg correction for multiple testing.
